# Identification of a Micropeptide Linked to Cancer Stem Cell Regulation and Chemoresistance

**DOI:** 10.1101/2023.03.14.532696

**Authors:** Achala Fernando, Chamikara Liyanage, Srilakshmi Srinivasan, Panchadsaram Janaththani, Jyotsna Batra

**Affiliations:** School of Biomedical Sciences, Faculty of Health, Queensland University of Technology, Brisbane, QLD 4059, Australia; The Centre for Genomics and Personalised Health, Queensland University of Technology, Brisbane, QLD 4059, Australia; Translational Research Institute, Queensland University of Technology, Brisbane, QLD 4012, Australia

**Keywords:** Micropeptides, stem cells, prostate cancer, IRX cluster, IRX4_PEP1, chemoresistance

## Abstract

Short open reading frames encoding micropeptides (miPEPs) less than 100 amino acids in length have recently emerged as important regulators of diverse biological functions. However, the functional role of cancer-specific miPEPs in cancer progression and therapeutic response remains largely unexplored. Genome-wide association studies have identified an association of Iroquois (IRX) clusters with multiple cancer risk. In this study, we identified 17 miPEPs generated from IRX clusters in prostate, breast, endometrial, and ovarian cancers using SWATH-MS/MS-based proteomic analysis. We found that *IRX4*-derived miPEP, IRX4_PEP1, promotes prostate cancer (PCa) cell proliferation, migration, and invasion by interacting with heterogeneous nuclear ribonucleoprotein K (hnRNPK). Overexpression of IRX4_PEP1 leads to dysregulation of stem cell pathways by co-interaction with Catenin beta-1 (CTNB1) and upregulation of prominent PCa stem markers, resulting in docetaxel resistance in PCa. IRX4_PEP1 expression is significantly upregulated in prostate tumour tissues compared to normal and is positively correlated with disease aggressiveness.

**SIGNIFICANCE:** Our findings highlight the critical role of IRX4_PEP1 in regulating PCa stem cells and chemotherapy resistance and suggest it as a novel therapeutic target for the treatment of PCa. Moreover, IRX4_PEP1 expression can serve as a novel potential diagnostic and prognostic biomarker for PCa.

## INTRODUCTION

In recent years, several unannotated mammalian short open reading frames (sORFs) have been discovered that have the potential to encode micropeptides (miPEPs) less than 100 amino acids in length (1,2). These sORFs can be located in various regions, including the 5′-untranslated region (5’-UTR), 3’-untranslated region (3’-UTR), non-coding RNAs and alternatively spliced transcripts (3,4). MiPEPs have been linked to various cancer hallmarks including neoplastic transformation, metastasis, chemo, and drug resistance (5-7). Despite their potential significance in cancer research, hormone-sensitive cancers such as prostate, breast, endometrial and ovarian cancers have been largely neglected in miPEP research. This is particularly relevant given that cancer stem-like cells (CSCs), which have been found to drive cancer initiation, relapse, metastasis, and therapy resistance in multiple cancers, including prostate cancer (PCa), have not been fully characterized in terms of miPEP regulation (8,9). Therefore, identification and profiling of miPEPs in clinical samples using the data-independent acquisition method of Sequential window acquisition of all theoretical mass spectra (SWATH– MS), holds promise for shedding light on the role of miPEPs in hormone-sensitive cancers and CSC regulation.

SWATH–MS is a data–independent acquisition method and has a high coverage of proteome with quantitatively and accurately (10). In SWATH–MS/MS, almost all peptides within a specified mass range are fragmented in an unbiased and systematic manner using windows of large precursor isolation, therefore it is a feasible method for identification and profiling of miPEPs in clinical samples (10-12).

The Iroquois (IRX) family, a subgroup of the homeobox family encode transcription factor proteins, have fundamental regulatory roles in patterning and differentiation in vertebrates (13). Although homeobox gene deregulation is strongly linked with many human malignancies, IRX family has not yet been fully functionally explored (8,14,15). The IRX family includes six genes, *IRX1, IRX2*, and *IRX4* (IRX cluster A) which are located in chromosome 5 and *IRX3, IRX5*, and *IRX6* (IRX cluster B) which are located in chromosome 16 (13,16). Genome-Wide Association Studies (GWAS) have recently identified several cancer risk-associated single nucleotide polymorphisms within the IRX clusters in patient cohorts pointing to their emerging role in carcinogenesis (17-24). The *5p15* locus bearing the IRX cluster A is associated with prostate, lung and gastric cancer risk (17,20,25-29) while the *16q12* locus bearing IRX cluster B is associated with breast, endometrial, and retinoblastoma cancer risk (30-32). However, the underlying mechanism of IRX cluster function in cancer progression and their ability to encode functional miPEPs remains a mystery, despite their importance in oncogenesis.

In this study, we utilized SWATH-MS PRIDE clinical data from prostate, breast, endometrial, and ovarian cancers to identify miPEPs generated by the IRX cluster. We compared the differential expression of these miPEPs across different cancer subtypes, grades, and stages, and identified IRX4_PEP1 as a promising candidate for functional characterization. Through overexpression and knockdown experiments, we found that IRX4_PEP1 significantly impacted PCa cell proliferation, migration, and invasion, and increased docetaxel resistance and stemness in PCa via interacting with specific protein partners. Further analysis showed that IRX4_PEP1 expression was significantly upregulated in tumour tissues compared to adjacent non-malignant tissues and correlated with PCa aggressiveness. Based on these findings, we propose that targeting IRX4_PEP1 may represent a novel therapeutic strategy for overcoming therapeutic resistance in PCa.

## RESULTS

### Identification of miPEPs generated from IRX clusters

SWATH-MS/MS raw data was obtained from PCa (PXD004691) (33), breast cancer (PXD014194) (34), endometrial cancer (PXD017217) and ovarian cancer (PXD010437) (35) from PRoteomics IDEntifications Database (PRIDE) database. A total of 17 miPEPs encoded by distinct *IRX* sORFs were identified in breast, endometrial, prostate and ovarian cancers using a proteogenomic pipeline explained in methods and Figure S1. A total of ten miPEPs were identified from IRX cluster A, three, four and three miPEPs from *IRX1, IRX2*, and *IRX4*, respectively and seven new miPEPs from IRX cluster B, five, one and one miPEPs from *IRX3, IRX5* and *IRX6*, respectively (Figure 1A). AUG was considered as the start codon if there was an upstream in-frame AUG codon. If no upstream AUG was available, an in-frame near-cognate non-AUG codon was selected as the start codon. About ∼29 % of miPEPs had canonical start codons (AUG) while ∼71% had near-cognate start codons (Figure 1B). The majority of the miPEPs (∼53%) were derived from predicted *IRX* transcripts/ spliced transcripts, about 24% from 3’UTRs and 5’UTRs (Figure 1C). From the total identified miPEPs, 82% of miPEPs were longer than 41 amino acids (Figure 1D). *In silico* CPPred-sORF analysis to predict the coding potential of 17 miPEPs sORFs based on SVM classifier and multiple sequence features suggested that 13/17 have a high coding potential above 51% and 4/17 have a low coding potential below 51% (Figure 1E; Table S1) (36).

**Figure 1.**
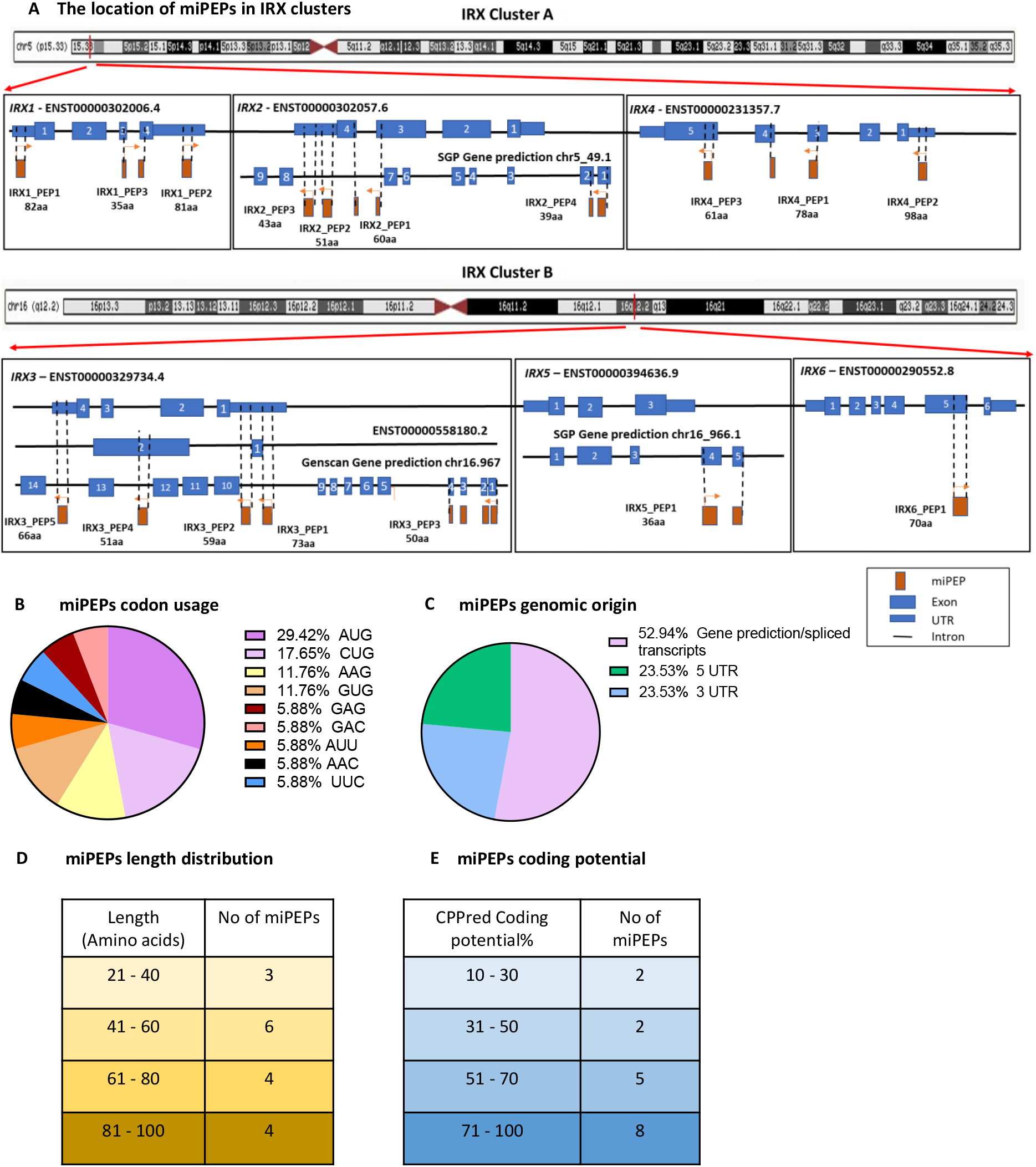
Proteomic identification of IRX cluster-encoded miPEPs in prostate, breast, endometrial and ovarian cancers. **(A)** The RNA map illustrating the alignment of sORFs that are translated into miPEPs in each IRX cluster A and B, including 5′UTR, 3′UTR, and predicted/ spliced transcripts. The blue boxes represent the annotated exons for each *IRX* transcript, the thin blue boxes represent UTRs and the orange boxes represent the miPEP and its alignment with the *IRX* sORFs. The arrows on the miPEP show the direction of the translation of sORFs. **(B)** The distribution of the start codon usage of miPEPs based on canonical and non-canonical start sites. **(C)** The distribution of miPEPs based on their genomic origin. **(D)** The distribution of miPEPs by their length in amino acids. **(E)** The predicted coding potential of miPEPs using CPPred-sORF software. See also Figure S1 and Table S1.

### Profiling of IRX cluster-derived miPEPs expression in patient samples

Eight, seven, six and three miPEPs were detected in the breast, endometrial, prostate and ovarian cancers (Figure 2A). No common peptides were detected for all cancers, however, IRX4_PEP3 was detected in all except ovarian cancer (Table S1). The miPEPs identified in breast cancer, IRX2_PEP2, IRX2_PEP3, IRX2_PEP4, IRX3_PEP1, IRX3_PEP2, IRX3_PEP5, IRX4_PEP3 and IRX5_PEP1 were quantified across the different cancer subtypes. Overall, the human epidermal growth factor Receptor positive (Her) subtype showed the highest expression of IRX miPEPs, while the triple-negative (TN) showed the lowest (Figure 2B). IRX3_PEP2 miPEP was significantly differentially expressed between ductal carcinoma in situ (DCIS), estrogen and progesterone receptor (ERPR) and Her positive subtypes (Figure S2A). The miPEPs identified in endometrial cancer, IRX1_PEP2, IRX3_PEP1, IRX3_PEP4, IRX4_PEP2, IRX4_PEP3, IRX5_PEP1 and IRX6_PEP1 were quantified across cancer grades (Figure 2C). A significantly lower expression of IRX5_PEP1 was observed in Grade 2 and Grade 3 compared to Grade 1 samples (Figure S2B).

**Figure 2.**
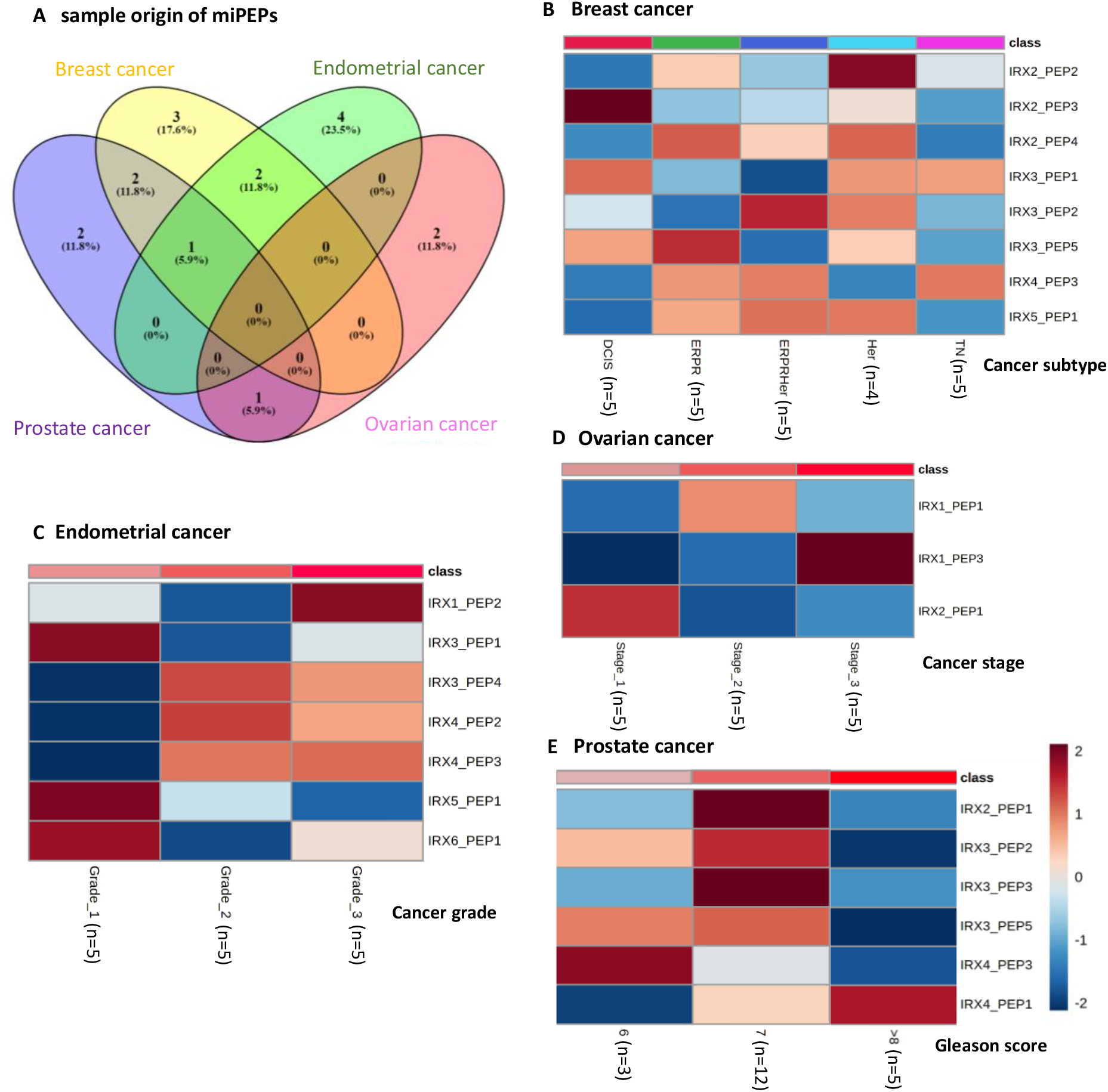
Quantification of miPEPs expression in breast, endometrial, ovarian and prostate cancer clinical specimens. **(A)** The Venn diagram represents the number of identified IRX miPEPs from SWATH-MS clinical datasets in PCa (PXD004691) (33), breast cancer (PXD014194) (34), endometrial cancer (PXD017217) and ovarian cancer (PXD010437) (35) and the number of overlapping miPEPs in these cancers. The heatmaps indicate the expression of identified IRX-miPEPs in **(B)** breast cancer by subtypes, Ductal carcinoma in situ (DCIS), Estrogen and Progesterone Receptor (ERPR) positive, human epidermal growth factor Receptor (Her) positive and ERPRHer positive and triple negative (TN) **(C)** endometrial cancer by cancer grades, Grade 1, 2 and 3 **(D)** ovarian cancer by cancer stages, Stage1, 2 and 3 **(E)** PCa by Gleason score, 6,7 and >8 of the clinical samples. The log2 transformed intensities of the miPEPs were z-scored before use in the heatmap. The number of clinical specimens used for each category has been shown in the brackets. The coloured scale bar for the intensity of miPEPs expression is shown. See also Figure S2.

The miPEPs identified in ovarian cancer, IRX1_PEP1, IRX1_PEP3, and IRX2_PEP1 were quantified across tumour stages where IRX1_PEP3 showed significant differential expression between stages 1 and 3 (Figure 2D, Figure S2C). The miPEPs identified in PCa, IRX2_PEP1, IRX3_PEP2, IRX3_PEP3, IRX3_PEP5, IRX4_PEP1, and IRX4_PEP3 were quantified across Gleason score (Figure 2E). IRX4_PEP3 showed a significant downregulation in Gleason score 6 to 7 and IRX4_PEP1 showed a significant upregulation with high Gleason score (Figure S2D). Overall, the differential expression of IRX miPEPs suggests that they are potential biomarkers to distinguish the subtypes of breast cancer, grades and stages of endometrial cancer, ovarian cancer and PCa.

### Overexpression and knockdown of IRX4_PEP1 impact PCa cell proliferation,migration and invasion

Since *IRX4*-derived IRX4_PEP1 miPEP expression is specific to PCa and is upregulated with high Gleason grade, IRX4_PEP1 was selected for further characterization. IRX4_PEP1 is 78 amino acids in length with an AUG canonical start codon and have a high CPPred-sORF coding potential of 91%. Our previous studies related to the *IRX4* locus have identified a potential transcript variant that can encode IRX4_PEP1 and have validated its expression at the protein level in PCa cell lines (18).

IRX4_PEP1 differs from the IRX4 transcription factor as it lacks DNA binding and transactivation domains coding regions, thus the secondary structure of IRX4_PEP1 was predicted using I-TASSER software (37). The results showed that the peptide has 2 helixes and coils with a relatively high confidence score (Figure 3A) for its amino acid sequence. The top predicted model for IRX4_PEP1 showed a C-score of -3.94 (Figure 3B). C -score is a confidence score normally in the range of (−5 to 2), where a C-score of higher value signifies a model with high confidence and vice-versa. Based on the extent of the inherent thermal mobility of residues in proteins, a normalized B-factor was predicted for the IRX4_PEP1, where negative values indicate high stability in the structure of the protein. As in Figure 3C, the first coil and the helix have shown negative values whereas most of the other residues show values around zero suggesting a stable nature for the IRX4_PEP1. According to SignalP -5.0 signal peptide prediction software, no signal peptides were predicted for IRX4_PEP1, suggesting its probability to be a non-secreted protein (38) (Figure S3A). Together *in silico* results suggests the stable nature of IRX4_PEP1 in cellular systems and support the importance of discovering its functional role.

**Figure 3.**
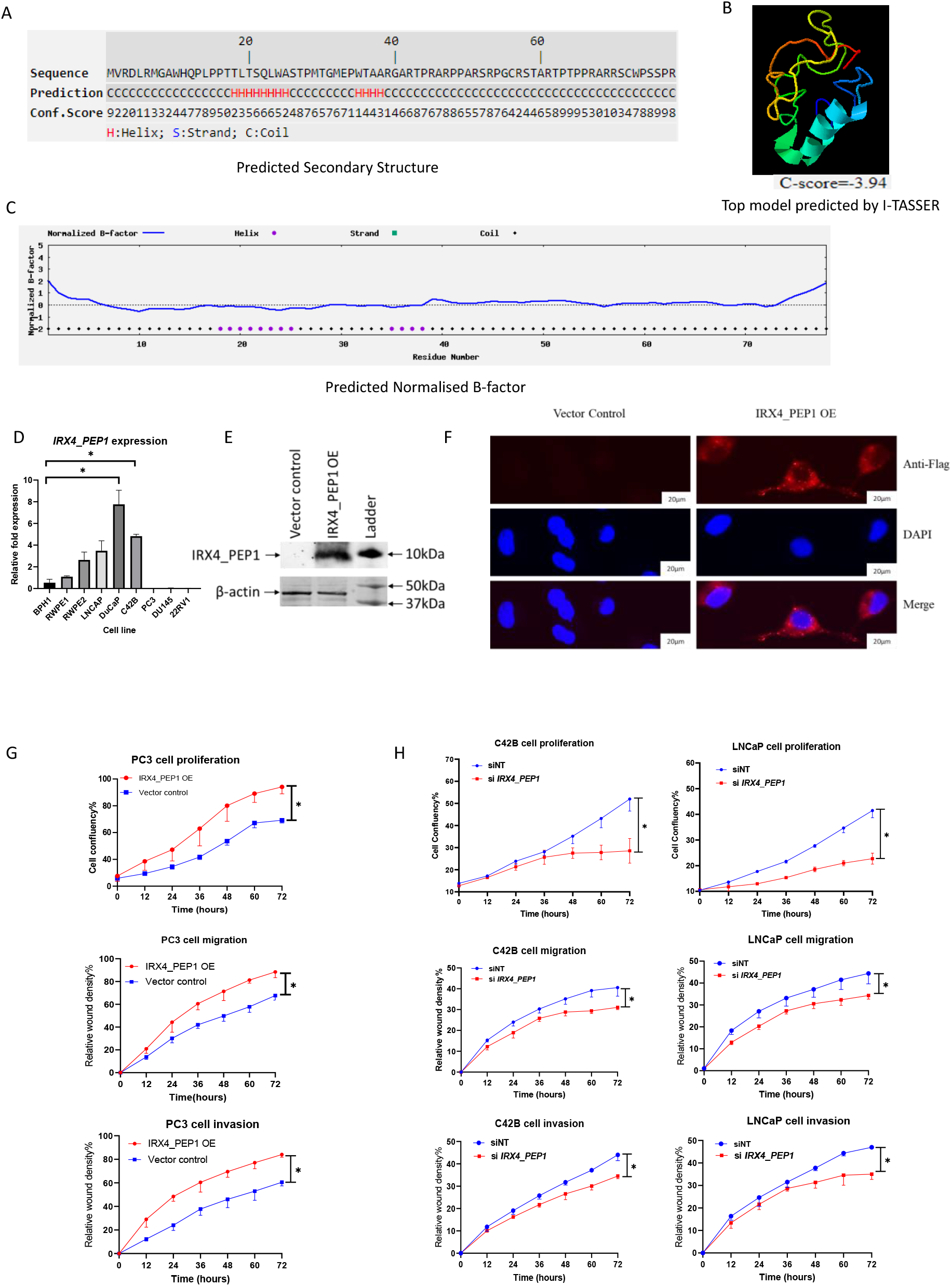
Overexpression (OE) and knockdown of IRX4_PEP1 impact PCa cell proliferation, migration, and invasion. **(A)** The predicted secondary structure for IRX4_PEP1 by I-Tasser H: Helix, S: Strand, C: Coil by I-Tasser. **(B)** The top predicted model for IRX4_PEP1(C-score= -3.94) by I-Tasser. **(C)** The predicted normalized B-factor for IRX4_PEP1 by I-Tasser. The B-factor profile in the figure corresponds to the normalized B-factor of the target protein, defined by B=(B’-u)/s, where B’ is the raw B-factor value, u and s are respectively the mean and standard deviation of the raw B-factors along the sequence. **(D)** The *IRX4_PEP1* RNA expression in a panel of PCa cell lines by qRT-PCR. The graph shows the relative quantification of *IRX4_PEP1* RNA expression compared to BPH1, benign PCa cell line. *RPL32* was used as the loading control (n=3 biological replicates, Mean±SD, Mann-Whitney test, *p<0.05) **(E)** The western blot with anti-Flag (DYK) antibody shows IRX4_PEP1 peptide expression (∼10kDa) in PC3 OE cells compared to vector control. β-actin was used as the loading control. **(F)** IRX4_PEP1 subcellular localisation validation with stable OE PC3 cells compared to vector control by an Immunofluorescence (IF) assay using anti-Flag (DYK) antibody **(G)** Incucyte proliferation, migration, and invasion assays with IRX4_PEP1 stable OE PC3 cells in comparison to vector control. **(H)** Incucyte proliferation, migration, and invasion assays with IRX4_PEP1 transient knockdown LNCaP and C42B cells in comparison to vector control. The images of the cells were taken in two-hour intervals for 72 hours and the confluency and/or relative wound density was analysed using the IncuCyte live-cell analysis system. (n=3 biological replicates, Mean±SD, Friedman test with Dunn’s multiple comparisons test, *p<0.05). See also Figure S3.

First, the expression analysis of *IRX4_PEP1* RNA across a panel of PCa cell lines identified DuCaP, an androgen-responsive cell line with the highest expression followed by C42B a CRPC cell line (Figure 3D). Androgen-responsive PCa cell lines, LNCaP and DuCaP showed upregulation of *IRX4_PEP1* compared to the androgen-independent cell lines, PC3 and DU145, suggesting an androgen-dependent regulation of IRX4_PEP1(Figure 3D). C42B, LNCaP (expressing moderate *IRX4_PEP1*) and PC3 cell lines (expressing low *IRX4_PEP1*) were shortlisted for additional functional studies using knockdown and OE models, respectively. Doxycycline inducible flag tagged-IRX4_PEP1 stable OE models in PC3 cells demonstrated a significant OE efficiency of *IRX4_PEP1* (over 200-fold) compared to the vector control (Figure S3B). The coding potential of the IRX4_PEP1 (∼10 kDa) was validated by a Western blot with an anti-flag (DYK) antibody (Figure 3E) and the densitometry analysis revealed a significant OE efficiency of IRX4_PEP1 (over 50-fold) compared to the vector control (Figure S3C). Immunofluorescence (IF) assay to validate the sub-cellular localization of IRX4_PEP1, suggested IRX4_PEP1 to predominantly localize to the cytoplasm (Figure 3F). Incucyte cell functional assays showed significantly higher proliferation, migration and invasion rates in IRX4_PEP1 OE PC3 cells compared to the vector control (Figure 3G). Significant knockdown efficiency >50% of *IRX4_PEP1* was achieved for both cell types using *IRX4_PEP1* specific siRNA compared to the non-targeting siRNA control (Figure S3D, E). Incucyte cell functional assays showed significantly lower proliferation, migration and invasion rates compared to the non-targeting siRNA control in both LNCaP and C42B on transient knockdown of *IRX4_PEP1*(Figure 3H).

### IRX4_PEP1 impacted stem cell regulatory pathways in PCa

The SWATH-MS proteomic profiling of IRX4_PEP1 OE PC3 cells identified differentially regulated 3,516 proteins between IRX4_PEP1 and vector control (Table S2). Ingenuity Pathway Analysis (IPA) revealed potential functional pathways associated with dysregulated proteins (Table S3). Interestingly, canonical pathways related to stem cell regulation were highlighted such as mTOR Signaling (p-value = 0.00023), GM-CSF Signaling (p-value = 0.00363), JAK/Stat signalling (p-value = 0.025) and embryonic stem cell pluripotency (p-value = 0.0331) (Figure 4A).

**Figure 4.**
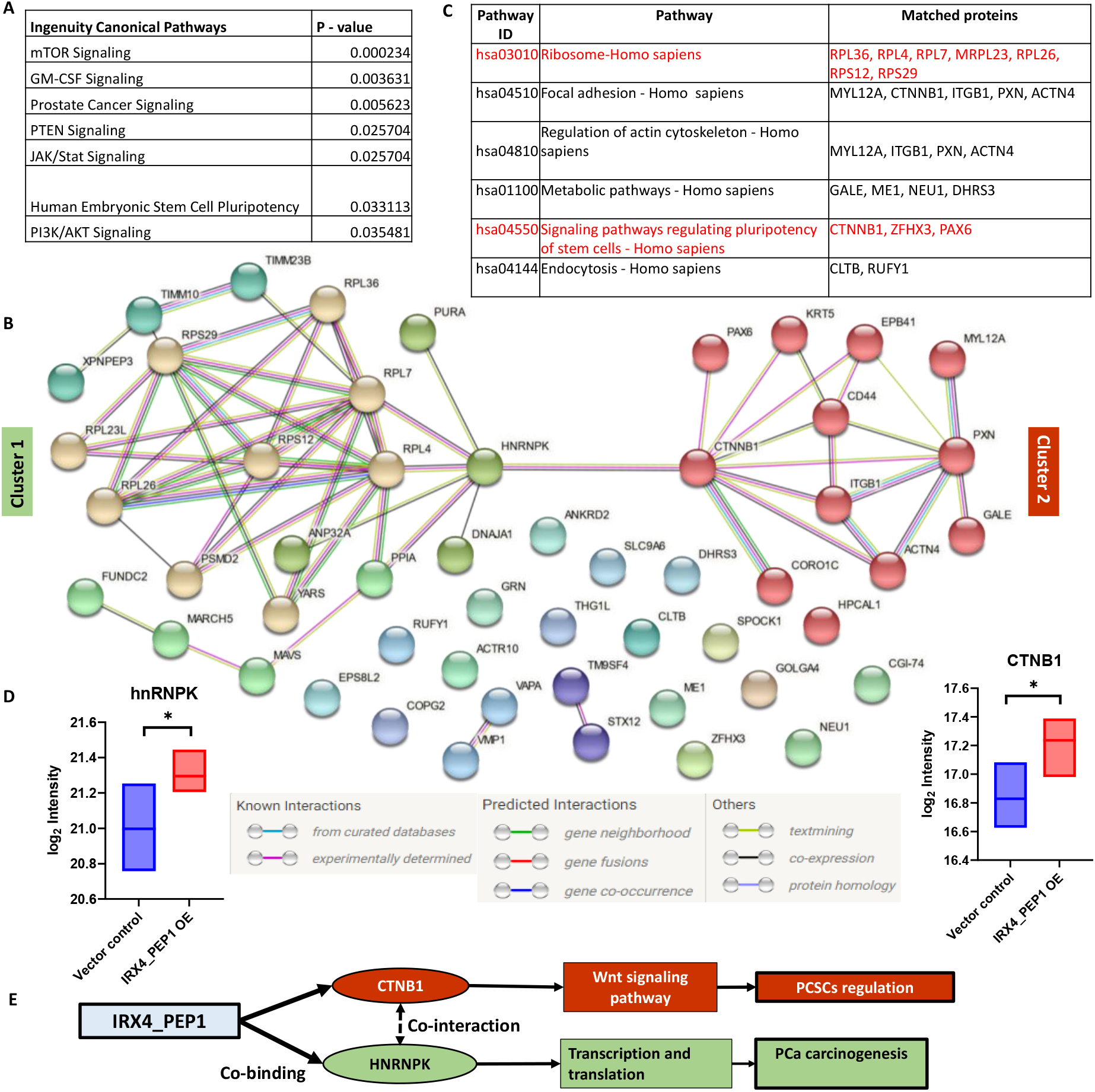
IRX4_PEP1 impacted stem cell regulatory pathways in PCa. **(A)** Top canonical pathways identified by whole proteomic dysregulated proteins with IRX4_PEP1 OE in PC3 cells by IPA pathway analysis. **(B)** The STRING protein analysis of the identified 52 potential co-interacting proteins of IRX4_PEP1 by Co-IP assay, two distinct clusters were observed, cluster 1 and cluster 2 as shown in distinct colours. **(C)** The KEGG pathway analysis of identified potential co-interacting proteins of IRX4_PEP1. The KEGG pathway ID, KEGG pathway, and the co-proteins mapped with KEGG pathway associated molecules have shown in table. **(D)** The bar graph shows the log2 normalized intensity of hnRNPK and CTNB1 proteins, the co-IP regulator proteins of two clusters in IRX4_PEP1 OE samples compared to vector control (n=3 biological replicates, *P<0.05, Student t-test). **(E)** The associated pathways related to co-proteins hnRNPK and CTNB1 in PCa according to previous literature (39-42). See also Figure S4.

Co-immunoprecipitation (Co-IP) analysis identified a total of 52 proteins that potentially interact with IRX4_PEP1(Table S4). Predicted STRING protein network analysis of the identified co-interacting proteins showed two main distinct protein clusters/networks, cluster 1 regulated by Heterogeneous nuclear ribonucleoprotein K (hnRNPK) and cluster 2 regulated by Catenin beta 1 protein (β catenin; CTNB1) (Figure 4B). Moreover, the CD44 protein, a prominent stem cell marker in PCa was identified as a co-interacting protein of IRX4_PEP1 (Figure 4B). Interestingly, KEGG signalling pathway analysis with identified co-interacting proteins of IRX4_PEP1 picked up pluripotency of stem cells regulating signalling pathways and ribosome-related pathways. The pathway list and the identified matched IRX4_PEP1 co-interacting proteins with each KEGG pathway proteins are shown in the table (Figure 4C, Figure S4). The expression of two main co-IP cluster regulator proteins, hnRNPK and CTNB1 were significantly upregulated with IRX4_PEP1 OE compared to vector control (Figure 4D), proving IRX4_PEP1 may bind to hnRNPK and CTNB1 proteins and subsequently upregulate their expression. Interestingly, CTNB1 has been identified as a stem cell regulator associated with the Wnt signalling pathway that drives prostate CSCs (PCSCs) (39) and hnRNPK, a prominent transcriptional and translational regulator, that drives PCa carcinogenesis, CRPC and stem cell regulation (Figure 4E) (40-42).

### IRX4_PEP1 induces stemness in PCa cells

To elucidate the rationale on PCSC regulation by IRX4_PEP1, a stem cell enrichment assay was carried out by single-cell seeding followed by CD44 staining according to a published protocol (43). IRX4_PEP1 stable OE PC3 cells were used, following 2 weeks of single-cell seeding, three types of colonies were detected that were characterized as holoclones, meroclones and paraclones (Figure 5A), of which holoclones are expected to have more stem-like features (43). A significant low percentage of holoclones (IRX4_PEP1 OE -17% and vector control -8%) and a higher percentage of non-holoclones, meroclones and paraclones (IRX4_PEP1 OE - 83% and vector control -92%) were detected from all the sustained clones indicating that stem cells are a rare population in PCa cells (Figure 5B). The cells from individual clones were sorted through flow cytometry following CD44 staining which showed enrichment of the holoclones with CD44^high^ stem-like cells (97.1%) compared to the non-holoclones (54.2%) (Figure 5C). Moreover, IRX4_PEP1 OE samples showed a higher percentage of CD44^high^ cells in both holoclones and non-holoclones than vector control suggesting a role in stemness upregulation by IRX4_PEP1 (Figure 5C). *IRX4_PEP1* mRNA levels in CD44 pre-sorted and post-sorted samples were measured compared to the vector control and confirmed the OE efficiency throughout the assay process (Figure 5D).

**Figure 5.**
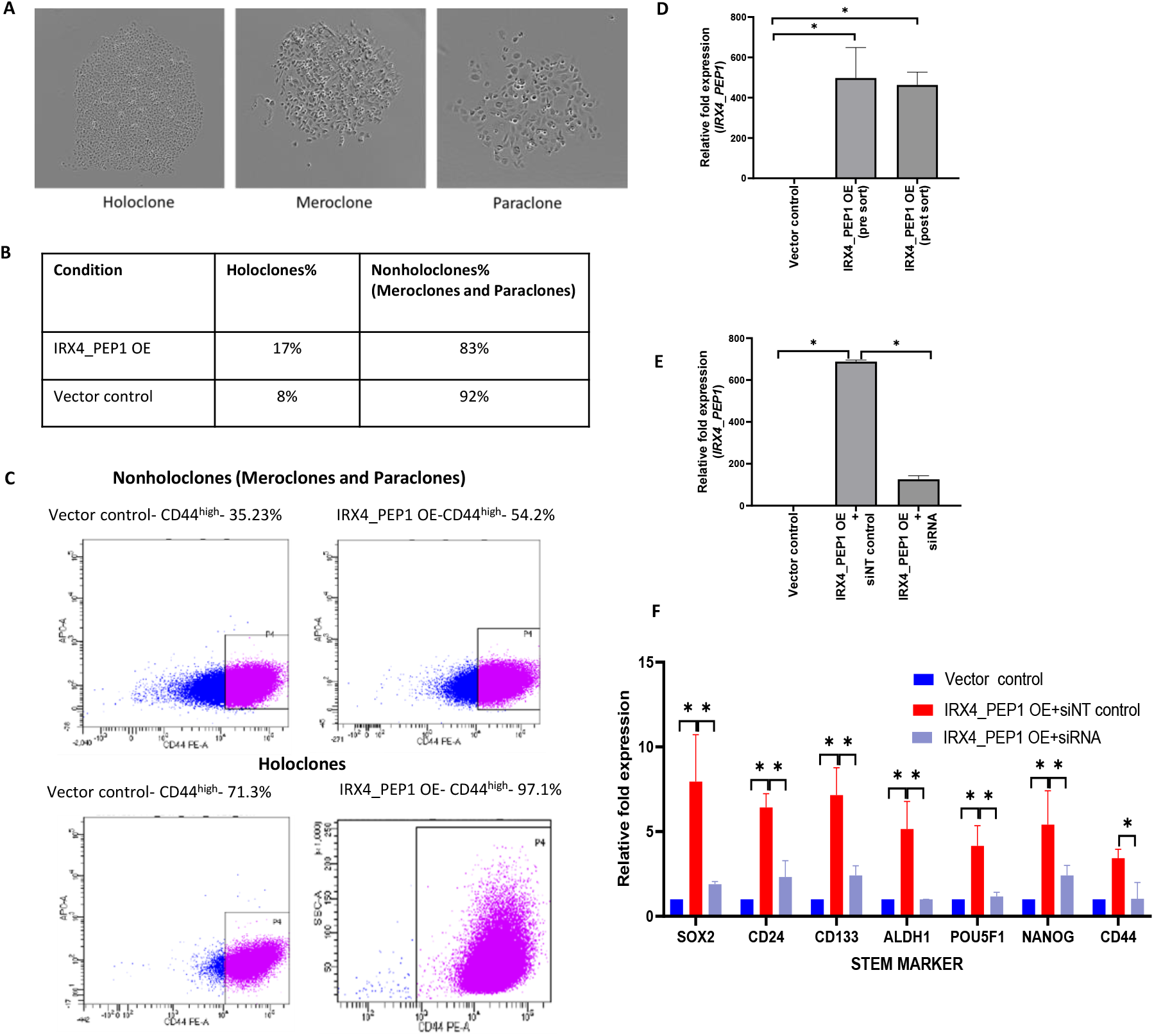
IRX4_PEP1 induces stemness in PCa cells. **(A)** Characterization of colonies following 2 weeks of single cell seeding; 3 colony types were found holoclones, meroclones and paraclones. **(B)** The percentage of colonies, holoclones and non-holoclones (meroclones and paraclones) detected in IRX4_PEP1 OE and vector control plates. **(C)** The percentage of CD44^high^ cells (pink) and CD44^low^ cells (blue) derived from holoclones and non-holoclones by flow cytometry analysis following CD44 staining. **(D)** The relative *IRX4_PEP1* RNA expression in doxycycline-treated IRX4_PEP1 OE PC3 cells compared to vector control in pre-and post-CD44 sorted samples (Mean±SD, *p<0.05, student t-test). **(E)** The RNA expression of *IRX4_PEP1* in doxycycline-treated, IRX4_PEP1 OE holoclones (dox+siNT) and IRX4_PEP1 OE PC3 cells treated with *IRX4_PEP1* siRNA (dox+siIRX4) (Mean±SD, *p<0.05, student t-test). **(F)** The expression of prominent stem cell markers in IRX4_PEP1 OE and knocked-down holoclones. *RPL32* has been used as a housekeeping gene (n=3 biological replicates, Mean±SD, *p<0.05, Mann-Whitney test).

Expression of prominent stem cell markers was measured in CD44^high^ stem-like IRX4_PEP1 OE PC3 holoclones and subsequently knocked down with the *IRX4_PEP1* siRNA. A significant OE and knockdown efficiency of *IRX4_PEP1* was observed in holoclones (Figure 5E). Interestingly, the expression of prominent stem cell markers, *SOX2, CD24, CD133, ALDH1, POU5F1* and *Nanog*, were elevated significantly in IRX4_PEP1 OE holoclones and downregulated significantly on knockdown with *IRX4_PEP1* siRNA (Figure 5F). The elevation of *CD44* marker expression was not significant compared to the vector control as IRX4_PEP1 OE holoclones were sorted based on high CD44 expression. However, a significant downregulation of CD44 expression was observed with *IRX4_PEP1* knockdown (Figure 5F) confirming that IRX4_PEP1 induces stemness in PCa cells.

### IRX4_PEP1 induced chemoresistance in PCa

Given the upregulated expression of *IRX4_PEP1* in the metastatic castration-resistant cell line C42B and its regulatory role in PCa stemness, it is hypothesized that IRX4_PEP1 may also contribute to chemoresistance. Therefore, the association between IRX4_PEP1 expression and docetaxel resistance was investigated. PC3, LNCaP and C42B cell lines were treated with docetaxel and IC50 values on cell survival were measured. Interestingly, PC3, which has the minimum expression of *IRX4_PEP1* showed the lowest IC50 value (0.437) while LNCaP and C42B which have relatively high expression of *IRX4_PEP1* showed higher IC50 values, 5.07, 2.65 respectively, suggesting an association of high *IRX4_PEP1* expression with longer cell survival (Figure 6A).

**Figure 6.**
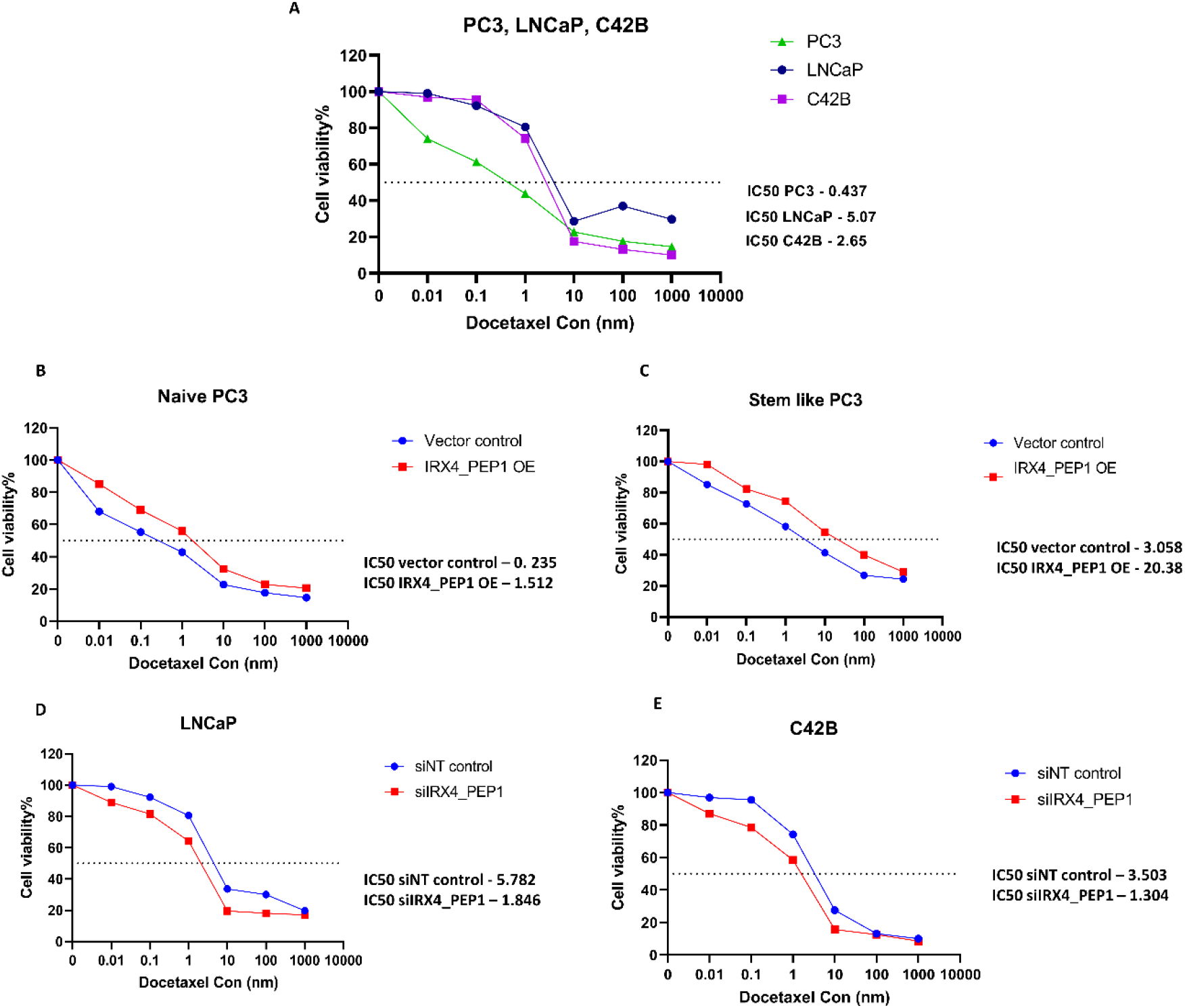
IRX4_PEP1 induced chemoresistance in PCa. **(A)** Dose response effects of docetaxel on PCa cell lines, PC3, LNCaP and C42B. **(B)** Dose response effects of docetaxel on IRX4_PEP1 OE naïve PC3 over vector control. **(C)** Dose response effects of docetaxel on CD44^high^ stem like IRX4_PEP1 OE PC3 over vector control. **(D)** Dose response effects of docetaxel on *IRX4_PEP1* transient knocked down LNCaP cells (siIRX4_PEP1) over non-targeting (siNT) control. **(E)** Dose response effects of docetaxel on IRX4_PEP1 transient knocked down C42B cells (siIRX4_PEP1) over non-targeting (siNT) control. 0.01, 0.1, 1, 10, 100, and 1000nM concentrations of docetaxel were used on prostate cancer cell lines, naïve PC3, CD44^high^ PC3, LNCaP and C42B over 48 hours. Normalised presto blue fluorescence absorbances were plotted for cell viability% and IC50 values for each cell line and treatment were presented (n=3 biological replicates).

IRX4_PEP1 associated docetaxel resistance was evaluated using IRX4_PEP1 OE naïve PC3 cells compared to vector control. Interestingly, IRX4_PEP1 OE naïve PC3 cells showed a higher percentage of viable cells in all docetaxel concentrations (IC50 - 1.512) compared to the parental naive PC3 (IC50 - 0.235) cells (Figure 6B). Moreover, on docetaxel treatment, CD44^high^ stem cell-like IRX4_PEP1 OE PC3 cells showed a higher cell survival (IC50 - 20.38) compared to vector control (IC50 – 3.058) suggesting that IRX4_PEP1 may induce docetaxel resistance in both naïve and stem like PCa cells (Figure 6C). It is interesting to note that stem-like PC3 cells showed a higher percentage cell survival (IC50-3.058) compared to the parental naive PC3 (IC50-0.235) (Figure 6B and C).

To validate the above results, LNCaP and C42B cell lines which endogenously express *IRX4_PEP1* were knocked down with *IRX4_PEP1* siRNA (siIRX4_PEP1) and treated with different concentrations of docetaxel. Lower percentage of viable cells were observed in both LNCaP and C42B knock down cell lines (IC50-1.846 and 1.304 respectively) compared to the siNT control (IC50-5.782 and 3.503) (Figure 6D and E) suggesting that, knockdown of *IRX4_PEP1* may reduce docetaxel resistance in PCa.

### IRX4_PEP1 expression is upregulated in PCa, CRPC and gradually increases with high Gleason score and high Gleason grade

The expression of IRX4_PEP1 was explored in PCa patient samples for its likely potential as a biomarker of PCa progression. The Cancer Genome Atlas Program (TCGA) PCa transcriptomic data analysis revealed that *IRX4_PEP1* expression is significantly increased in PCa tumour samples (n=49) compared to non-malignant prostatic samples (n=49) (Figure 7A). PRIDE reprocessing of SWATH-MS dataset (PXD004691)(33) showed the significant upregulation of IRX4_PEP1 in PCa tumour samples (n=20) compared to non-malignant samples (n=20) (Figure 7B). Reprocessing the public SWATH-MS dataset (PASS01126)(44) has indicated that IRX4_PEP1 expression was significantly increased in metastatic CRPC samples (n=11) compared to localised primary PCa samples (n=9) (Figure 7C) further confirming IRX-PEP1 induced treatment resistance.

**Figure 7.**
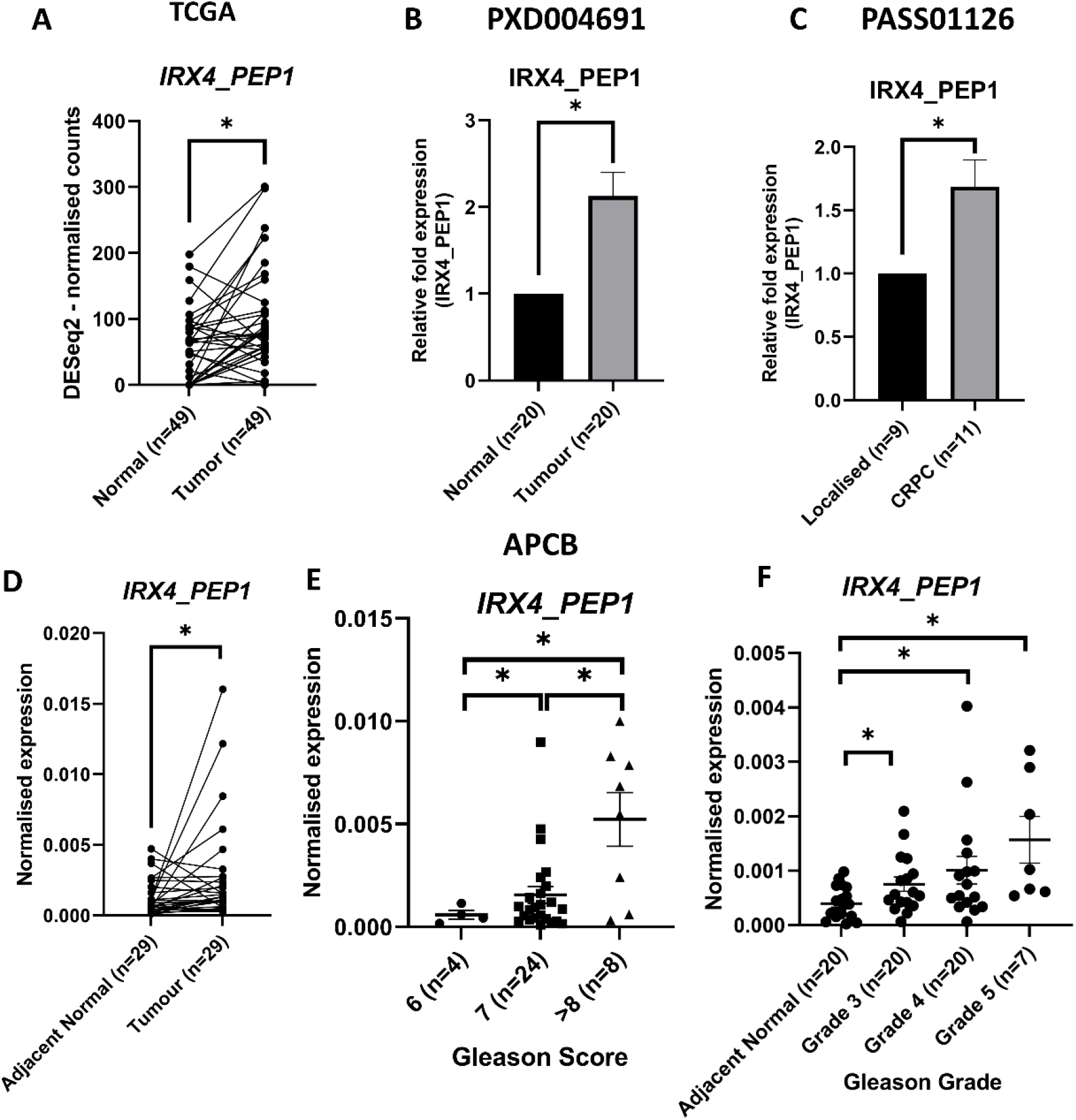
IRX4_PEP1 expression is upregulated in PCa, CRPC clinical samples and correlates with high PCa Gleason score and Gleason grade. **(A)** *IRX4_PEP1* mRNA expression in PCa TCGA samples (n=49) and non-malignant prostate tissue samples (n=49). *RPL32* was used as the housekeeping gene. Each patient is represented by a single dot in the graph (n=3 technical replicates, paired t-test *p<0.05). **(B)** IRX4_PEP1 relative protein expression in PCa samples (n=20) compared to non-malignant prostate tissue samples (n=20) (PRIDE PXD004691) Mean±SD, student t-test *p<0.05. **(C)** IRX4_PEP1 relative protein expression in localised primary PCa samples (n=9) compared to CRPC samples (n=11) (PASS01126) Mean±SD, student t-test *p<0.05. **(D)** *IRX4_PEP1* mRNA expression in APCB samples (n=29) and adjacent non-malignant prostate tissue samples (n=29). *RPL32* was used as the housekeeping gene. Each patient is represented by a single dot in the graph (n=3 technical replicates, paired t-test *p<0.05). **(E)** *IRX4_PEP1* mRNA expression in APCB patient samples across different Gleason scores. *RPL32* was used as the housekeeping gene (n=3 technical replicates, Mean±SD, Mann-Whitney test *p<0.05). **(F)** *IRX4_PEP1* mRNA expression in APCB patient samples across different Gleason grades. *RPL32* was used as the housekeeping gene (n=3 technical replicates, Mean±SD, Mann-Whitney test *p<0.05).

Furthermore, analysing tumour samples of 29 Australian Prostate Cancer BioResource (APCB) patients and their adjacent non-malignant tissues, demonstrated upregulated *IRX4_PEP1* expression in tumour samples (Figure 7D). Interestingly, *IRX4_PEP1* expression gradually increased with high Gleason score (Figure 7E) and positively correlated with higher Gleason grades (Figure 7F) implying its potential as a diagnostic biomarker to stratify prostate tumour from normal epithelial tissues as well as the potential to be a prognostic biomarker in PCa progression.

## DISCUSSION

The sORFs that generate miPEPs can play significant functions in crucial biological processes and cancer hallmarks (5-7). GWAS and functional follow-up studies reported an association of IRX clusters with cancer risk. However, the underlying mechanisms of IRX clusters in cancer functions remain a mystery. We hypothesised, the IRX cluster can encode functional miPEPs and these miPEPs can have a role in cancer progression. We utilised a computational-proteomic workflow to identify IRX cluster encoded miPEPs by reprocessing SWATH-MS/MS PRIDE raw data of hormone-sensitive cancers, as IRXs seem to be hormone-regulated according to our previous findings (18). SWATH-MS/MS analysis is a robust, highly reproducible label-free quantification method that has been used for clinical sample analysis in many studies (11,12,45). In this study, we identified 17 miPEPs that are cancer-specific and are common to several cancers irrespective of heterogeneity between samples suggestive of their high conservation. Majority of the identified miPEPs expression varied between cancer stages, grades and subtypes suggesting their biomarker potential to stratify different stages and subtypes of cancer. Majority of miPEPs were derived from alternatively spliced transcripts implying alternative splicing is a great source of sORFs and miPEPs, as also determined by previous studies (1,7). IRX4_PEP1 is a miPEP derived from a transcript variant of *IRX4* with a high coding potential and an AUG canonical start codon. Its expression is specific to PCa and increases with advanced PCa, suggesting its significance in this cancer type. In our previous study, a transcript variant of *IRX4* that encode this miPEP was detected in both PCa cell lines and clinical samples (18). As a result, IRX4_PEP1 was chosen for further functional characterization.

Our *in-silico* modelling of IRX4_PEP1 suggests a stable nature of this miPEP in cellular systems. The endogenous expression of *IRX4_PEP1* was identified in many PCa cell lines at RNA level, however the expression at the protein level was confirmed with a Flag antibody as IRX4_PEP1 specific antibody is not available. Our functional results suggest an oncogenic role of IRX4_PEP1 in PCa as its overexpression significantly increased PCa cell proliferation, migration and invasion, while transient knockdown of *IRX4_PEP1* showed opposite effects. Whole proteomic and IPA pathway analysis of proteins dysregulated by IRX4_PEP1 OE compared to vector control revealed that majority of the pathways impacted were stem cell signalling pathways. The above findings were further validated with Co_IP results which suggests that the potential interacting proteins of IRX4_PEP1 are also members of several stem cell signalling pathways.

Interestingly, Catenin beta-1 (CTNB1) was identified as one of the potential interacting protein of IRX4_PEP1 as well as a main regulator of one of the STRING protein cluster. CTNB1 is a core component of the Wnt/CTNB1 signalling pathway which regulates stemness in PCa (46). Moreover, upregulated expression of CTNB1 was detected with IRX4_PEP1 OE compared to the vector control. IRX4_PEP1 binding to CTNB1 may upregulate CTNB1 expression in PCa and may subsequently activate the Wnt pathway that regulates stemness in PCa. In line with this result, a previous study has detected that LINC00689 and miR-496 binding to CTNB1 activates the Wnt pathway in PCa (47). Moreover, CTNB1 activation drives androgen receptor (AR)-independent CRPC progression (48), which is in line with our data, showing high expression of IRX4_PEP1 in CRPC samples compared to localised PCa.

PCSCs are responsible for driving the progression of advanced PCa and CRPC (49). However, the isolation of PCSCs from in-house PCa cells for stem cell research remains a significant challenge. We developed an effective model for enriching PCSCs from in-house PCa cells adapting a previously published protocol (50-52). Our model enabled us to detect different colony morphologies including round, tightly packed cell colonies (Holoclones) and irregularly shaped colonies consisting of larger, loosely packed cells (Meroclone and paraclones). Among these, holoclones are enriched with more CSCs and have the highest proliferative capacity (50-52). The holoclones were a better model for CSCs studies in PCa, which we utilised in our study together with high CD44 expression, the most common marker for isolating PCSCs (51,53). Our results confirmed that IRX4_PEP1 induced PCa stemness, while upregulating expression of prominent PCa stem cell markers.

In addition, our results indicate that, IRX4_PEP1 binds to heterogeneous nuclear ribonucleoprotein K (hnRNPK), one of the major proteins involved in mRNA translation (40) and may upregulate the expression of hnRNPK affecting the ribosomal protein (PRPLs), regulating the translational network in PCa. HnRNPK upregulation at RNA and protein levels in PCa has been identified previously, however the direct association with PCa cell proliferation has not been explored (54). Instead, hnRNPK together with Purα has been found to bind to the *AR* and the hnRNPK–AR-related signature contributes to CRPC progression (41,55). The hnRNPK activates c-Myc and its downstream targets genes, leading to cell cycle progression that drives gastric and breast cancer cell proliferation (56,57), suggesting a similar mechanism may apply in PCa together with IRX4_PEP1. In addition, a physical interaction of hnRNPK with CTNB1 has been reported in neuroblastoma where hnRNPK binding increases the stability and transactivation of CTNB1 (58). In PCa, this interaction has not been reported, however our results suggest the possibility of an interaction between hnRNPK and CTNB1 and this complex may promote tumour growth and stemness upregulation in PCa. However, our results merely are not sufficient to conclude that the PCa stem cell regulation and high cell proliferation are associated with individual functions of CTNB1 and hnRNPK, respectively or from the complex together. High proliferative capacities of CSCs compared to naïve cancer cells were evident in previous literature. For example, a stem cell line CSC480 developed from a normal colon cancer cell line, demonstrated enhanced proliferative capacity compared to the colon cancer cell line, SW480. The proliferation marker, Ki67, is expressed by CSC480 cells at a 160% level higher than of SW480 cells (59). Another example is that injection of acute myeloid leukemia CSCs into immunosuppressed mice could initiate tumour formation by 30-to 100-fold higher than normal cells indicating significant self-renewal and proliferative capability of stem cells (60).

CSCs-induced chemoresistance has been reported in many cases (61,62). IRX4_PEP1 seems to reduce sensitivity to docetaxel in both stem-like cells and naïve PCa cells that may be influenced by upregulated stem markers. Moreover, an increased docetaxel resistance in stem-like cells was observed than parental PCa cells, consistent with the previous studies (63). Knock down of *IRX4_PEP1*, successfully restored sensitivity to docetaxel in PCa cells suggesting IRX4_PEP1 may be a novel therapeutic target in chemo resistant PCa.

IRX4_PEP1 is significantly upregulated in PCa patient samples and showed continuous upregulation with a high Gleason score and high Gleason grade, suggesting the use of IRX4_PEP1 expression for more accurate prediction on disease progression. The Gleason score is used widely and is calculated as the sum of the two most prevalent Gleason grades, primary (most prevalent pattern) and secondary (any amount of a worse pattern) (64). During PCa diagnosis, detecting the actual prognosis of the treatment is vital as it can direct personalised as well as more patient-oriented therapies. *IRX4_PEP1* mRNA expression may be useful in combination with Gleason grade and Gleason score for more accurate prediction of PCa progression, planning the treatment course to achieve a better outcome for patients. Validation of *IRX4_PEP1* expression in a large patient cohort would strengthen the IRX4_PEP1’s promise as a potential prognostic marker.

The miPEP identification and their expression analysis largely vary on diverse proteomic analysis methods, varied sample processing, and different bioinformatic pipelines and may be related to heterogeneity in cancer subtypes. Therefore, the clinical value of miPEPs primarily relies on establishing more systematic functional investigation rather than relying on robust analytical tools. Herein, our results identified functional association of IRX4_PEP1 in PCa, however further mechanistic assays are required to understand IRX4_PEP1-influenced complete cell programming.

In summary, it is evident that hidden small ORFs in IRX cluster has the potential translation capability to encode functional miPEPs. *IRX4* derived newly identified miPEP, IRX4_PEP1 plays an oncogenic role in PCa by upregulating stemness and chemoresistance. Targeting IRX4_PEP1 could open novel therapeutic opportunities to halt tumour progression, stem cell associated relapse and chemoresistance in PCa. Moreover, IRX4_PEP1 may be a potential diagnostic and prognostic biomarker for PCa. Taken together, IRX4_PEP1 supports the phenomenon that discovering miPEPs linked to tumorigenesis, and their functional characterisation will open new avenues in cancer therapy. Moreover, the result hints that IRX miPEPs-based biomarkers may not be limited to hormone-sensitive cancers but rather apply to many other cancers with diverse tissue origins.

## METHODS

### Cell Culture

The cell line-based studies are approved by the Queensland University of Technology (QUT) ethics committee (QUT Ethics approval number: 1500001082). A panel of PCa cell lines (LNCaP, DuCaP, C42B, DU145, PC3, 22RV1, RWPE2) and benign prostate cell lines (BPH1, RWPE1) were purchased from the American Type Culture Collection (ATCC, Manassas, VA, USA). The cell lines were grown in RPMI1640 (1X) with no phenol red (Life Technologies, Grand Island, NY, USA) supplemented with 5% fetal bovine serum except for RWPE-1 and RWPE-2 cell lines which were grown in a keratinocyte serum-free medium supplemented with 5 ng/mL recombinant human epidermal growth factor (EGF) and 50 μg/mL bovine pituitary extract (Gibco™, Invitrogen, Carlsbad, CA, USA). The cell lines were tested for short tandem repeat (STR) profiling and tested negative for mycoplasma.

### Retrieving sequences for alternative *IRX4* transcripts from Proteomics UCSC genome browser

We identified a total of 67 human *IRX* transcripts newly from UCSC genome browser derived from alternative splicing, gene predictions and UTRs. A database was created from hypothetical peptide sequences of 865 different ORFs that were translated from 67 transcripts. All the frames with canonical and non-canonical start sites of hypothetical peptide sequences were selected. CPPred-sORF tool was utilised to predict the coding potential of identified sORFs (36).

### Retrieving mass-spectrometry raw files from PRIDE database

PRIDE database containing the DIA SWTH-MS/MS raw files were systematically mined to retrieve raw files related to prostate, breast and endometrial cancer tissue samples (PXD004691 (33), PXD014194 (34), PXD017217 with. wiff SWATH files respectively) and ovarian cancer PXD010437 (35)with thermos. raw files). For prostate cancer, 40 LC-MS/MS raw files representing 20 prostate cancer and their matched normal FFPE tissue samples were obtained. For breast cancer patient analysis, 24 LC-MS/MS raw files were retrieved of 5 in each category of DCIS, ERPR, ERPRHer, TN and 4 in Her sub-cancer categories. For endometrial cancer, 15 LC-MS/MS raw files of FFPE tumour tissues were retrieved of 5 in each category representing 3 cancer grades and for ovarian cancer 15 LC-MS/MS raw files of FFPE tumour tissues were retrieved of 5 in each category representing 3 cancer stages. Localised primary PCa (n=9) and CRPC (n=11) SWATH-MS data was obtained from the Peptide Atlas repository (PASS01126) (44).

### SWATH-MS/MS Data Analysis of PRIDE Proteomic Datasets

A pipeline was developed to identify and quantify miPEPs that are being translated by IRX sORFs using SWATH-MS/MS datasets of clinical samples. The spectral library was created using data-dependent acquisitions (DDA). All the data-dependent acquisitions were imported into ProteinPilot™ software (Version 5.0.1, AB SCIEX) and searched using the Paragon™ algorithm against the FASTA database consisting of newly identified IRX miPEPs sequences merged with the UniProt human reference database and contaminant proteins. The following search parameters were used: Sample type: Identification; Cys Alkylation: Iodoacetamide; Digestion: Trypsin; Instrument: TripleTOF 5600+; Species: None; Search effort: Thorough ID; Results Quality: Detected protein threshold [Unused ProtScore (Conf)] ≥ 0.05 with False discovery rate (FDR) The generated ion library was imported into PeakView® SWATH micro app (Version 2.1, AB SCIEX) along with the SWATH files (. wiff format files). 1%-FDR and >90% confidence with 2 minimum peptides and 3 minimum transitions was set to identify IRX miPEPs. Created ion library was cleaned and saved in text format using the iSwathX tool (Version 2.0). The curated ion library was imported into Skyline software (Version 4.2) along with the DIA data (Thermo.raw format) for the DDA/DIA peptide search using 0.90 cut-off score and the following parameters: precursor charges: 2, 3; mass tolerance: 10 ppm; mass tolerance: 0.05 Da; enzyme: trypsin, missed cleavages: 2. and miPEP quantification was performed using MStats R-based statistical tool (Version 2.0). Identified peptides was blasted against the human proteome to remove peptides matched with annotated human protein/peptide sequences using NCBI blast. Finally, 17 previously unannotated miPEPs were shortlisted for further analysis of their expression based on tumour subtypes, stages, and grades.

### Tissue collection from APCB patient samples

Human ethic approval was obtained for APCB sample collection (QUT ethics approval no-1000001165). APCB clinical and their adjacent normal FFPE clinical samples were serially sectioned (20 μm) and stained with hematoxylin and eosin. Microdissection was performed manually on the tumour Gleason grades marked FFPE areas using a sterile injection needle and deparaffinized using deparaffinization solution (Qiagen, Catalog number – 19093). RNA extraction was performed using the miRNeasy FFPE kit (Qiagen, Catalog number – 217504) according to the manufacturer’s instructions. RNA quality and quantity were measured using NanoDropTM1000 (Thermo Scientific, Biolab, Scoresby, VIC, Australia). RNA from 47 APCB patient samples were used in the analysis. In addition, matching 20 adjacent normal non-malignant tissue samples were utilised.

### RNA Isolation, cDNA Synthesis, Relative Quantification by Real-Time Quantitative RT-PCR (qRT-PCR)

Total RNA was extracted from PCa cells using the Isolate II RNA Mini Kit (Bioline, London, UK) and was reverse transcribed to cDNA using the SensiFastTM cDNA synthesis kit (Bioline, GmbH, Luckenwalde, Germany). The primers for qRT-PCR were designed using the NCBI tool Primer-BLAST–NCBI–NIH software. All the primer sequences are given in Table S5. RNA quality and quantity were measured using NanoDrop™1000 Quantitative RT-PCR was performed using the ViiA7 Real-Time PCR system (Applied Biosystems, Foster City, CA, USA). Each reaction contained 1X final concentration of SYBR Green PCR Master Mix 2X (Applied Biosystems, Foster City, CA, USA), 50 nM forward and reverse primer, 2.0 μL of diluted cDNA (1:5) and nuclease-free water at a final volume of 8 μL. The cycling parameters were 95 °C for 10 min, 40 cycles of 95 °C for 15 s and 60 °C for 1 min followed by a dissociation step. All the CT values were normalized to the expression of the housekeeping gene *RPL32* (ΔCT). Relative expression compared to control was performed by the comparative CT (ΔΔCT) method.

### IRX4_PEP1 stable OE cell establishment

The full length of *IRX4_PEP1* was PCR amplified and cloned into the pDONR223 vector (Addgene) with the DYKDDDDK Flag tag by the BP recombination reaction protocol (Gateway® technology, Invitrogen) using BP clonase enzyme mix (Invitrogen, cat no 11789020). Then the insert was cloned into a doxycycline-inducible lentiviral vector, pINDUCER21(ORF-EG) (Addgene) by the LR recombination reaction protocol (Gateway® technology (Invitrogen) using the LR clonase enzyme mix (Invitrogen, cat no 11791019). The inserts were confirmed by sequencing and proceeded to the lentiviral transfection. 12 μL X-treme GENE HP DNA transfection Reagent with 1.8 μg of pCMV-Δ8.2R, 0.2 μg of pCMV-VSVG and 2.0 μg of pDNA (plasmid of interest) was prepared and was then dripped onto the 293T cells and was incubated overnight at 37°C. The viral supernatant was harvested at 48 hours. The target PC3 cells were infected with the viral supernatant and protamine sulphate (8 μg/ml). The cell sorting was done based on GFP fluorescence. The IRX4_PEP1 OE was achieved with the treatment of 0.25ug doxycycline hylate (Sigma-Aldrich D9891-5G) and no doxycycline treated cells were used as the vector control.

### IRX4_PEP1 transient knockdown assay

IRX4_PEP1 transfected PC3 cells were transiently transfected with 10nM siIRX4_PEP1 siRNA (Invitrogen by Life Technologies, cat no # 1299001) using Lipofectamine® RNAiMAX transfection Reagent (Invitrogen, cat no -13778150) in the presence and absence of doxycycline. After 48 hours of transfection at 37 °C, total RNA was isolated. The negative control was non-targeting siRNA (Silencer select no 1 siRNA, cat no – 4390843, Ambion).

### Western blot analysis

The doxycycline-treated and non-treated whole cell lysate was isolated with RIPA buffer (50mM Tri-HCl pH 7.5, 150mM NaCl, 1% SDS, 1% Triton X-100 and 1x Protease Inhibitor Cocktail) and the western blot was carried out (Odyssey® system) with Anti-Flag (DYKDDDDK Tag (D6W5B) Rabbit mAb#14793 antibody (Cell Signaling). Beta-actin (PA1-183, Thermo Fisher) was used as the loading control. Briefly, the protein concentrations were measured using a standard BCA assay with Pierce™ BCA Protein Assay Kit (Sigma). 30 μg of proteins was loaded into a pre-cast gel (Novex™ 16% Tricine Gels, 1.0 mm, 10 well cat no EC6695BOX) and followed manufacturer instructions.

### Incucyte® Cell Proliferation assay

The cells were seeded in a 96-well plate with a density of 3000 cells/well following the siRNA transfection or doxycycline treatment. The two images per well were taken every two hours in the Incucyte (essenbioscience.com) for three consecutive days and cell confluence was plotted. The experiment was performed with 3 biological and 3 technical replicates with a non-targeting control/vector control.

### Incucyte Wound Scratch assays

#### Migration/Invasion

The cells were seeded in a poly-L-Ornithine (Sigma-Aldrich, Catalog number - P4957) coated 96-well ImageLock plate (Essen BioScience, Catalog number - 4379) with a density of 50,000 cells/well. The next day, the cells were treated with 10μg/ml mitomycin C (Sigma-Aldrich, Catalog number – M4287) for 1 hour to prevent proliferation. After washing with media, the uniform wounds were created in all wells using the WoundMakerTM followed by the siRNA transfection or doxycycline treatment. The two images per well were taken every two hours for three consecutive days and the wound closure/relative wound density was plotted. The experiment was performed with 3 biological and 3 technical replicates with a non-targeting control/vector control. For invasion, an additional 2% Matrigel layer was added to the wells.

#### Immunofluorescence assay (IF)

To confirm the sub-cellular localisation of IRX4_PEP1 miPEP, the IRX4_PEP1 stable OE cells were fixed with 4% PFA and permeabilized with Triton X-100. Then the cells were treated with Anti-Flag antibody (DYKDDDDK Tag (D6W5B) Rabbit mAb#14793 antibody (Cell Signaling) overnight at 37 °C and incubated with anti-Rabbit IgG Secondary Antibody coupled with Alexa Fluor® 568 conjugate (red). The nuclei were counter stained with 4′,6-diamidino-2-phenylindole (DAPI) (blue) and imaged (40X) with an Olympus Inverted Fluorescence microscope.

#### Proteomic analysis of IRX4_PEP1 OE PC3 cells

IRX4_PEP1 transfected PC3 cells were used for whole proteome analysis to identify deregulated proteins by IRX4_PEP1 OE. Protein sample preparation, SWATH-MS/MS analysis, data analysis and IPA and KEGG pathway analysis were performed according to the protocol/method published in our previous study (12). The proteins with fold change ≤ -1.5 or ≥ 1.5 and the p-value < 0.05 were considered differentially regulated and used for further analysis.

#### Co-Immunoprecipitation (Co-IP) assay

IRX4_PEP1 transfected PC3 cell protein lysate was pre-cleared with 25 μL of SureBeads TM Protein G magnetic beads (BIO-RAD, Catalog number – 1614023) for 1 hour at 4 ºC and the beads were removed three times using a magnetic stand. The extract was then incubated with 5 μg of Anti-FLAG Antibody (DYKDDDDK Tag (D6W5B) Rabbit mAb#14793 antibody (Cell Signaling) and 50 μL of protein G beads at 4 ºC overnight. Beads were washed with ice-cold PBS three times and the proteins were eluted with 100 μL nuclear extraction buffer by boiling the samples at 95 ºC for 5 minutes. Then, the bound proteins which eluted were analysed by mass spectrometry at the Central Analytical Research Facility at QUT, using SWATH-MS/MS protocol published in our previous study (12).

#### Stem cell enrichment in IRX4_PEP1 transfected PC3 cell lines

IRX4_PEP1 transfected PC3 cells were collected and seeded in 96-well plates as one cell/well by using a single seeding facility in flow cytometry, at Translational Research Institute. Three replicate 96 well plates in each doxycycline and no doxycycline conditions were prepared. Wells with only one cell was selected and marked. Two weeks later, morphological classification of single cell clones was carried out, holoclones and non-holoclones were divided and sorted by flow cytometry for stem-like cells following CD44 staining with CD44 antibody (Miltenyi Biotec Cat no 130-095-180). Sorted CD44^high^ holoclones were used to isolate RNA to quantify stem cell marker expression.

#### RNA seq analysis of TCGA patient clinical samples

The RNA-seq data (bam format) of 49 PCa patients of TCGA was used in the study (“The Cancer Genome Atlas Program,” 2021). RASflow, HISAT2 and Deseq2 software were used for exon-specific RNA-seq analysis and data normalization (65,66).

#### PrestoBlue cell viability assay

IRX4_PEP1 transfected PC3 cells were cultured and treated with 0.01,0.1, 1, 10, 100, 1000nM docetaxel and 0.25ug doxycycline in a 96-well plate. After 48 hours of treatment, 10 μL of warmed cell viability reagent (Invitrogen™ catalogue number: A13261) was added to the 90 μL culture medium and the covered plate was incubated for 40min at 37 °C in a cell culture incubator. Following incubation, the fluorescence excitation readings were taken at a wavelength of 560 nm (the excitation range is 540–570 nm) and an emission of 590 nm (emission range is 580–610–nm). The viability counts were normalized to no doxycycline vector control. For knockdown assays, following siRNA transfection, different docetaxel concentrations were added and incubated for 48 hours. The viability counts were normalized to siNT control.

### Statistical analysis

All statistical data were analysed by GraphPad Prism 9.0.0 (121). The comparison was analysed by the student t test /Mann-Whiney test (two groups) and Friedman test with Dunn’s multiple comparisons (more than two groups). The results were considered statistically significant * if p < 0.05 at a 95% confidence interval. All the experiments were performed in 3 technical and 3 biological replicates. Exported SWATH-MS/MS peak area values were statistically analysed using the MetaboAnalyst 5.0 online tool. First, peak area values of each protein, obtained from the three replicates were averaged, normalized (based on the sum) and log_2_ transformed.

## Author Contributions

Conceptualization, J.B.; methodology, A.F., P.J., C.L., S.S; investigation, A.F, J.B; Statistical analysis A.F., C.L.; writing, original draft preparation, A.F., J.B.; writing-review and editing, all authors supervision, P.J. S.S., and J.B.; project administration, J.B.; funding acquisition, J.B. All authors have read and agreed to the published version of the manuscript.

## Acknowledgments

The authors acknowledge Adil Malik and Afshin Moradi for their contribution in the data analysis. Pawel Sadowski and Raj Gupta are acknowledged for carrying out the LC-MS/MS analysis at the Central Analytical Research Facility (CARF), operated by the Institute for Future Environments at QUT.

**The authors declare no competing interests**.

## Supplemental figure Legends

**Figure S1. Workflow to identify and quantify micro peptides in hormone-related cancers**.

The first step was to retrieve the nucleotide sequences of IRX novel ORFs from the UCSC genome browser followed by an *in-silico* translation to obtain the hypothetical peptide sequences and assembled a FASTA database with peptide sequences. The SWATH-MS/MS raw files originating from prostate, breast, endometrial and ovarian cancer patient tissue samples retrieved from the PRIDE database were subjected to ProteinPilot software for re-processing. Custom built FASTA database was used for peptide spectrum match (PSM) and identified unique peptides of IRXs (<100 amino acids) and quantified using PeakView and Skyline software. NCBI blast using UniProtKB databse was performed for homology detection with annotated Uniprot proteins and one identified miPEP was selected for further functional analysis in PCa. This figure is related to main **Figure 1**.

**Figure S2. Expression of significantly differentially expressed IRX-miPEPs in hormone-sensitive cancers based on the subtypes, grades and Gleason Score**.

**(A)** breast cancer by subtypes. **(B)** Endometrial cancer by cancer grades. **(C)** Ovarian cancer by cancer stages. **(D)** Prostate cancer by Gleason score. The normalised log2 intensities of the miPEPs were used to plot and the number of patients has been shown in the brackets. Mean±SD, *P<0.05. This figure is related to main **Figure 2**.

**Figure S3. Stable overexpression and Transient knockdown efficiency of IRX4_PEP1 at RNA and protein level**.

**(A)** According to SignalP - 5.0 signal peptide prediction software no signal peptide was predicted for IRX4_PEP1 miPEP (right), the figure has shown in comparison to PSA (left) which has a signal peptide peak. **(B)** The OE efficiency of IRX4_PEP1 RNA and **(C)** protein in doxycycline treated PC3 cells compared to vector control. *RPL32* has been used as a housekeeping gene. **(D)** IRX4_PEP1 transient knockdown efficiency in C42B and **(E)** LNCaP cells compared to nontargeting control (siNT). *RPL32* expression was used as the housekeeping control (n=3 biological replicates, Mean±SD, *p<0.05). This figure is related to main **Figure 3**.

**Figure S4. KEGG pathway analysis with potential co-partner proteins of IRX4_PEP1 identified by CO-IP assay**.

**(A)** The signalling pathway regulating pluripotency of stem cells and **(B)** ribosomal pathway that were mapped with KEGG analysis of co-partner proteins of IRX4_PEP1 has shown with the matched proteins highlighted. This figure is related to main **Figure 4**.

## Supplemental Table titles

Table S1. Identification and characterization of novel miPEPs generated from IRX cluster.

Table S2. Differentially regulated proteins with IRX4_PEP1 overexpression vs vector control.

Table S3. IPA pathway analysis with differentially regulated proteins with IRX4_PEP1 overexpression vs vector control.

Table S4. The potentially interacting proteins with IRX4_PEP1 miPEP.

Table S5. Primer and siRNA sequences were used in the study.

## REFERENCES

1. Slavoff SA, Mitchell AJ, Schwaid AG, Cabili MN, Ma J, Levin JZ, et al. Peptidomic discovery of short open reading frame-encoded peptides in human cells. Nat Chem Biol 2013;9(1):59–64 doi 10.1038/nchembio.1120.

2. Anderson DM, Anderson KM, Chang CL, Makarewich CA, Nelson BR, McAnally JR, et al. A micropeptide encoded by a putative long noncoding RNA regulates muscle performance. Cell 2015;160(4):595–606 doi 10.1016/j.cell.2015.01.009.

3. Dhamija S, Menon MB. Non-coding transcript variants of protein-coding genes - what are they good for? RNA Biol 2018;15(8):1025–31 doi 10.1080/15476286.2018.1511675.

4. Sousa ME, Farkas MH. Micropeptide. Plos Genet 2018;14(12):e1007764 doi 10.1371/journal.pgen.1007764.

5. Sun L, Wang W, Han C, Huang W, Sun Y, Fang K, et al. The oncomicropeptide APPLE promotes hematopoietic malignancy by enhancing translation initiation. Mol Cell 2021;81(21):4493–508 e9 doi 10.1016/j.molcel.2021.08.033.

6. Xiao MH, Lin YF, Xie PP, Chen HX, Deng JW, Zhang W, et al. Downregulation of a mitochondrial micropeptide, MPM, promotes hepatoma metastasis by enhancing mitochondrial complex I activity. Mol Ther 2022;30(2):714–25 doi 10.1016/j.ymthe.2021.08.032.

7. Ye M, Zhang J, Wei M, Liu B, Dong K. Emerging role of long noncoding RNA-encoded micropeptides in cancer. Cancer Cell Int 2020;20:506 doi 10.1186/s12935-020-01589-x.

8. Hanahan D. Hallmarks of Cancer: New Dimensions. Cancer Discov 2022;12(1):31–46 doi 10.1158/2159-8290.CD-21-1059.

9. Qiu ZX, Zhao LJ, Shen JZ, Liang ZY, Wu QL, Yang KL, et al. Transcription Elongation Machinery Is a Druggable Dependency and Potentiates Immunotherapy in Glioblastoma Stem Cells. Cancer Discov 2022;12(2):502–21 doi 10.1158/2159-8290.Cd-20-1848.

10. Ludwig C, Gillet L, Rosenberger G, Amon S, Collins BC, Aebersold R. Data-independent acquisition-based SWATH-MS for quantitative proteomics: a tutorial. Mol Syst Biol 2018;14(8):e8126 doi 10.15252/msb.20178126.

11. Narasimhan M, Kannan S, Chawade A, Bhattacharjee A, Govekar R. Clinical biomarker discovery by SWATH-MS based label-free quantitative proteomics: impact of criteria for identification of differentiators and data normalization method. J Transl Med 2019;17(1):184 doi 10.1186/s12967-019-1937-9.

12. Liyanage C, Malik A, Abeysinghe P, Clements J, Batra J. SWATH-MS Based Proteomic Profiling of Prostate Cancer Cells Reveals Adaptive Molecular Mechanisms in Response to Anti-Androgen Therapy. Cancers (Basel) 2021;13(4) doi 10.3390/cancers13040715.

13. Cavodeassi F, Modolell J, Gomez-Skarmeta JL. The Iroquois family of genes: from body building to neural patterning. Development 2001;128(15):2847–55.

14. Haria D, Naora H. Homeobox Gene Deregulation: Impact on the Hallmarks of Cancer. Cancer Hallm 2013;1(2-3):67–76 doi 10.1166/ch.2013.1007.

15. Wang XQD, Fan D, Han Q, Liu Y, Miao H, Wang X, et al. Mutant NPM1 Hijacks Transcriptional Hubs to Maintain Pathogenic Gene Programs in Acute Myeloid Leukemia. Cancer Discov 2023;13(3):724–45 doi 10.1158/2159-8290.CD-22-0424.

16. Peters T, Dildrop R, Ausmeier K, Ruther U. Organization of mouse Iroquois homeobox genes in two clusters suggests a conserved regulation and function in vertebrate development. Genome Res 2000;10(10):1453–62 doi 10.1101/gr.144100.

17. Dong J, Cheng Y, Zhu M, Wen Y, Wang C, Wang Y, et al. Fine mapping of chromosome 5p15.33 identifies novel lung cancer susceptibility loci in Han Chinese. Int J Cancer 2017;141(3):447–56 doi 10.1002/ijc.30702.

18. Fernando A, Liyanage C, Moradi A, Janaththani P, Batra J. Identification and Characterization of Alternatively Spliced Transcript Isoforms of IRX4 in Prostate Cancer. Genes (Basel) 2021;12(5) doi 10.3390/genes12050615.

19. Guo X, Liu W, Pan Y, Ni P, Ji J, Guo L, et al. Homeobox gene IRX1 is a tumor suppressor gene in gastric carcinoma. Oncogene 2010;29(27):3908–20 doi 10.1038/onc.2010.143.

20. Janaththani P. The role of GWAS identified 5p15 locus in prostate cancer risk and progression: Queensland University of Technology; 2019.

21. Nguyen HH, Takata R, Akamatsu S, Shigemizu D, Tsunoda T, Furihata M, et al. IRX4 at 5p15 suppresses prostate cancer growth through the interaction with vitamin D receptor, conferring prostate cancer susceptibility. Hum Mol Genet 2012;21(9):2076–85 doi 10.1093/hmg/dds025.

22. Udler MS, Ahmed S, Healey CS, Meyer K, Struewing J, Maranian M, et al. Fine scale mapping of the breast cancer 16q12 locus. Hum Mol Genet 2010;19(12):2507–15 doi 10.1093/hmg/ddq122.

23. Wu D, Li Z, Zhao S, Yang B, Liu Z. Downregulated microRNA-150 upregulates IRX1 to depress proliferation, migration, and invasion, but boost apoptosis of gastric cancer cells. Iubmb Life 2020;72(3):476–91 doi 10.1002/iub.2214.

24. Zhu Q, Wu Y, Yang M, Wang Z, Zhang H, Jiang X, et al. IRX5 promotes colorectal cancer metastasis by negatively regulating the core components of the RHOA pathway. Mol Carcinog 2019;58(11):2065–76 doi 10.1002/mc.23098.

25. Abate-Shen C. Deregulated homeobox gene expression in cancer: cause or consequence? Nat Rev Cancer 2002;2(10):777–85 doi 10.1038/nrc907.

26. Du JB, Xu YC, Dai JC, Ren CL, Zhu C, Dai NB, et al. Genetic variants at 5p15 are associated with risk and early onset of gastric cancer in Chinese populations. Carcinogenesis 2013;34(11):2539–42 doi 10.1093/carcin/bgt259.

27. McKay JD, Hung RJ, Gaborieau V, Boffetta P, Chabrier A, Byrnes G, et al. Lung cancer susceptibility locus at 5p15.33. Nat Genet 2008;40(12):1404–6 doi 10.1038/ng.254.

28. Petersen GM, Amundadottir L, Fuchs CS, Kraft P, Stolzenberg-Solomon RZ, Jacobs KB, et al. A genome-wide association study identifies pancreatic cancer susceptibility loci on chromosomes 13q22.1, 1q32.1 and 5p15.33. Nat Genet 2010;42(3):224–8 doi 10.1038/ng.522.

29. Zanetti KA, Wang Z, Aldrich M, Amos CI, Blot WJ, Bowman ED, et al. Genome-wide association study confirms lung cancer susceptibility loci on chromosomes 5p15 and 15q25 in an African-American population. Lung Cancer 2016;98:33–42 doi 10.1016/j.lungcan.2016.05.008.

30. Long JR, Cai QY, Shu XO, Qu SM, Li C, Zheng Y, et al. Identification of a Functional Genetic Variant at 16q12.1 for Breast Cancer Risk: Results from the Asia Breast Cancer Consortium. Plos Genet 2010;6(6) doi ARTN e100100210.1371/journal.pgen.1001002.

31. Susini T, Baldi F, Howard CM, Baldi A, Taddei G, Massi D, et al. Expression of the retinoblastoma-related gene Rb2/p130 correlates with clinical outcome in endometrial. Journal of Clinical Oncology 1998;16(3):1085–93 doi Doi 10.1200/Jco.1998.16.3.1085.

32. Priya K, Jada SR, Quah BL, Quah TC, Lai PS. High incidence of allelic loss at 16q12.2 region spanning RBL2/p130 gene in retinoblastoma. Cancer Biol Ther 2009;8(8):714–7 doi DOI 10.4161/cbt.8.8.7921.

33. Zhu Y, Weiss T, Zhang Q, Sun R, Wang B, Yi X, et al. High-throughput proteomic analysis of FFPE tissue samples facilitates tumor stratification. Mol Oncol 2019;13(11):2305–28 doi 10.1002/1878-0261.12570.

34. Valo I, Raro P, Boissard A, Maarouf A, Jezequel P, Verriele V, et al. OLFM4 Expression in Ductal Carcinoma In Situ and in Invasive Breast Cancer Cohorts by a SWATH-Based Proteomic Approach. Proteomics 2019;19(21-22):e1800446 doi 10.1002/pmic.201800446.

35. Thomas SN, Friedrich B, Schnaubelt M, Chan DW, Zhang H, Aebersold R. Orthogonal Proteomic Platforms and Their Implications for the Stable Classification of High-Grade Serous Ovarian Cancer Subtypes. iScience 2020;23(6):101079 doi 10.1016/j.isci.2020.101079.

36. Tong X, Liu S. CPPred: coding potential prediction based on the global description of RNA sequence. Nucleic Acids Res 2019;47(8):e43 doi 10.1093/nar/gkz087.

37. Roy A, Kucukural A, Zhang Y. I-TASSER: a unified platform for automated protein structure and function prediction. Nat Protoc 2010;5(4):725–38 doi 10.1038/nprot.2010.5.

38. Nielsen H, Tsirigos KD, Brunak S, von Heijne G. A Brief History of Protein Sorting Prediction. Protein J 2019;38(3):200–16 doi 10.1007/s10930-019-09838-3.

39. Qin W, Zheng Y, Qian BZ, Zhao M. Prostate Cancer Stem Cells and Nanotechnology: A Focus on Wnt Signaling. Front Pharmacol 2017;8:153 doi 10.3389/fphar.2017.00153.

40. Barboro P, Repaci E, Rubagotti A, Salvi S, Boccardo S, Spina B, et al. Heterogeneous nuclear ribonucleoprotein K: altered pattern of expression associated with diagnosis and prognosis of prostate cancer. Brit J Cancer 2009;100(10):1608–16 doi 10.1038/sj.bjc.6605057.

41. Capaia M, Granata I, Guarracino M, Petretto A, Inglese E, Cattrini C, et al. A hnRNP K-AR-Related Signature Reflects Progression toward Castration-Resistant Prostate Cancer. International Journal of Molecular Sciences 2018;19(7) doi ARTN 192010.3390/ijms19071920.

42. Xie W, Zhu HC, Zhao M, Wang L, Li SS, Zhao C, et al. Crucial roles of different RNA-binding hnRNP proteins in Stem Cells. Int J Biol Sci 2021;17(3):807–17 doi 10.7150/ijbs.55120.

43. Locke M, Heywood M, Fawell S, Mackenzie IC. Retention of intrinsic stem cell hierarchies in carcinoma-derived cell lines. Cancer Res 2005;65(19):8944–50 doi 10.1158/0008-5472.CAN-05-0931.

44. Latonen L, Afyounian E, Jylha A, Nattinen J, Aapola U, Annala M, et al. Integrative proteomics in prostate cancer uncovers robustness against genomic and transcriptomic aberrations during disease progression. Nat Commun 2018;9(1):1176 doi 10.1038/s41467-018-03573-6.

45. Singh AN, Sharma N. Quantitative SWATH-Based Proteomic Profiling for Identification of Mechanism-Driven Diagnostic Biomarkers Conferring in the Progression of Metastatic Prostate Cancer. Front Oncol 2020;10:493 doi 10.3389/fonc.2020.00493.

46. Schneider JA, Logan SK. Revisiting the role of Wnt/beta-catenin signaling in prostate cancer. Mol Cell Endocrinol 2018;462(Pt A):3–8 doi 10.1016/j.mce.2017.02.008.

47. Meng L, Li Z, Chen Y, Liu D, Liu Z. LINC00689 promotes prostate cancer progression via regulating miR-496/CTNNB1 to activate Wnt pathway. Cancer Cell Int 2020;20:215 doi 10.1186/s12935-020-01280-1.

48. Patel R, Brzezinska EA, Repiscak P, Ahmad I, Mui E, Gao M, et al. Activation of beta-Catenin Cooperates with Loss of Pten to Drive AR-Independent Castration-Resistant Prostate Cancer. Cancer Res 2020;80(3):576–90 doi 10.1158/0008-5472.CAN-19-1684.

49. Ojo D, Lin X, Wong N, Gu Y, Tang D. Prostate Cancer Stem-like Cells Contribute to the Development of Castration-Resistant Prostate Cancer. Cancers (Basel) 2015;7(4):2290–308 doi 10.3390/cancers7040890.

50. Beaver CM, Ahmed A, Masters JR. Clonogenicity: holoclones and meroclones contain stem cells. PLoS One 2014;9(2):e89834 doi 10.1371/journal.pone.0089834.

51. Duan JJ, Qiu W, Xu SL, Wang B, Ye XZ, Ping YF, et al. Strategies for isolating and enriching cancer stem cells: well begun is half done. Stem Cells Dev 2013;22(16):2221–39 doi 10.1089/scd.2012.0613.

52. Flynn L, Barr MP, Baird AM, Smyth P, Casey OM, Blackshields G, et al. Prostate cancer-derived holoclones: a novel and effective model for evaluating cancer stemness. Sci Rep 2020;10(1):11329 doi 10.1038/s41598-020-68187-9.

53. Skvortsov S, Skvortsova, II, Tang DG, Dubrovska A. Concise Review: Prostate Cancer Stem Cells: Current Understanding. Stem Cells 2018;36(10):1457–74 doi 10.1002/stem.2859.

54. Nagano K, Masters JR, Akpan A, Yang A, Corless S, Wood C, et al. Differential protein synthesis and expression levels in normal and neoplastic human prostate cells and their regulation by type I and II interferons. Oncogene 2004;23(9):1693–703 doi 10.1038/sj.onc.1207297.

55. Wang LG, Johnson EM, Kinoshita Y, Babb JS, Buckley MT, Liebes LF, et al. Androgen receptor overexpression in prostate cancer linked to Pur alpha loss from a novel repressor complex. Cancer Res 2008;68(8):2678–88 doi 10.1158/0008-5472.CAN-07-6017.

56. Mandal M, Vadlamudi R, Nguyen D, Wang RA, Costa L, Bagheri-Yarmand R, et al. Growth factors regulate heterogeneous nuclear ribonucleoprotein K expression and function. Journal of Biological Chemistry 2001;276(13):9699–704 doi DOI 10.1074/jbc.M008514200.

57. Peng WZ, Liu JX, Li CF, Ma R, Jie JZ. hnRNPK promotes gastric tumorigenesis through regulating CD44E alternative splicing. Cancer Cell International 2019;19(1) doi ARTN 33510.1186/s12935-019-1020-x.

58. Li D, Wang XJ, Mei H, Fang EH, Ye L, Song HJ, et al. Long Noncoding RNA pancEts-1 Promotes Neuroblastoma Progression through hnRNPK-Mediated beta-Catenin Stabilization. Cancer Research 2018;78(5):1169–83 doi 10.1158/0008-5472.Can-17-2295.

59. Alowaidi F, Hashimi SM, Alqurashi N, Alhulais R, Ivanovski S, Bellette B, et al. Assessing stemness and proliferation properties of the newly established colon cancer ‘stem’ cell line, CSC480 and novel approaches to identify dormant cancer cells. Oncol Rep 2018;39(6):2881–91 doi 10.3892/or.2018.6392.

60. Bonnet D, Dick JE. Human acute myeloid leukemia is organized as a hierarchy that originates from a primitive hematopoietic cell. Nat Med 1997;3(7):730–7 doi DOI 10.1038/nm0797-730.

61. Chandrasekar T, Yang JC, Gao AC, Evans CP. Mechanisms of resistance in castration-resistant prostate cancer (CRPC). Transl Androl Urol 2015;4(3):365–80 doi 10.3978/j.issn.2223-4683.2015.05.02.

62. de Morree ES, Vogelzang NJ, Petrylak DP, Budnik N, Wiechno PJ, Sternberg CN, et al. Association of Survival Benefit With Docetaxel in Prostate Cancer and Total Number of Cycles Administered: A Post Hoc Analysis of the Mainsail Study. JAMA Oncol 2017;3(1):68–75 doi 10.1001/jamaoncol.2016.3000.

63. Zhang X, Zhang S, Liu Y, Liu J, Ma Y, Zhu Y, et al. Effects of the combination of RAD001 and docetaxel on breast cancer stem cells. Eur J Cancer 2012;48(10):1581–92 doi 10.1016/j.ejca.2012.02.053.

64. Amin MB, Greene FL, Edge SB, Compton CC, Gershenwald JE, Brookland RK, et al. The Eighth Edition AJCC Cancer Staging Manual: Continuing to build a bridge from a population-based to a more “personalized” approach to cancer staging. CA Cancer J Clin 2017;67(2):93–9 doi 10.3322/caac.21388.

65. Kim D, Paggi JM, Park C, Bennett C, Salzberg SL. Graph-based genome alignment and genotyping with HISAT2 and HISAT-genotype. Nat Biotechnol 2019;37(8):907–15 doi 10.1038/s41587-019-0201-4.

66. Zhang X, Jonassen I. RASflow: an RNA-Seq analysis workflow with Snakemake. BMC Bioinformatics 2020;21(1):110 doi 10.1186/s12859-020-3433-x.

