## Supplementary figures and images for "Identification of a Micropeptide Linked to Cancer Stem Cell Regulation and Chemoresistance"

### Supplemental Figure 1

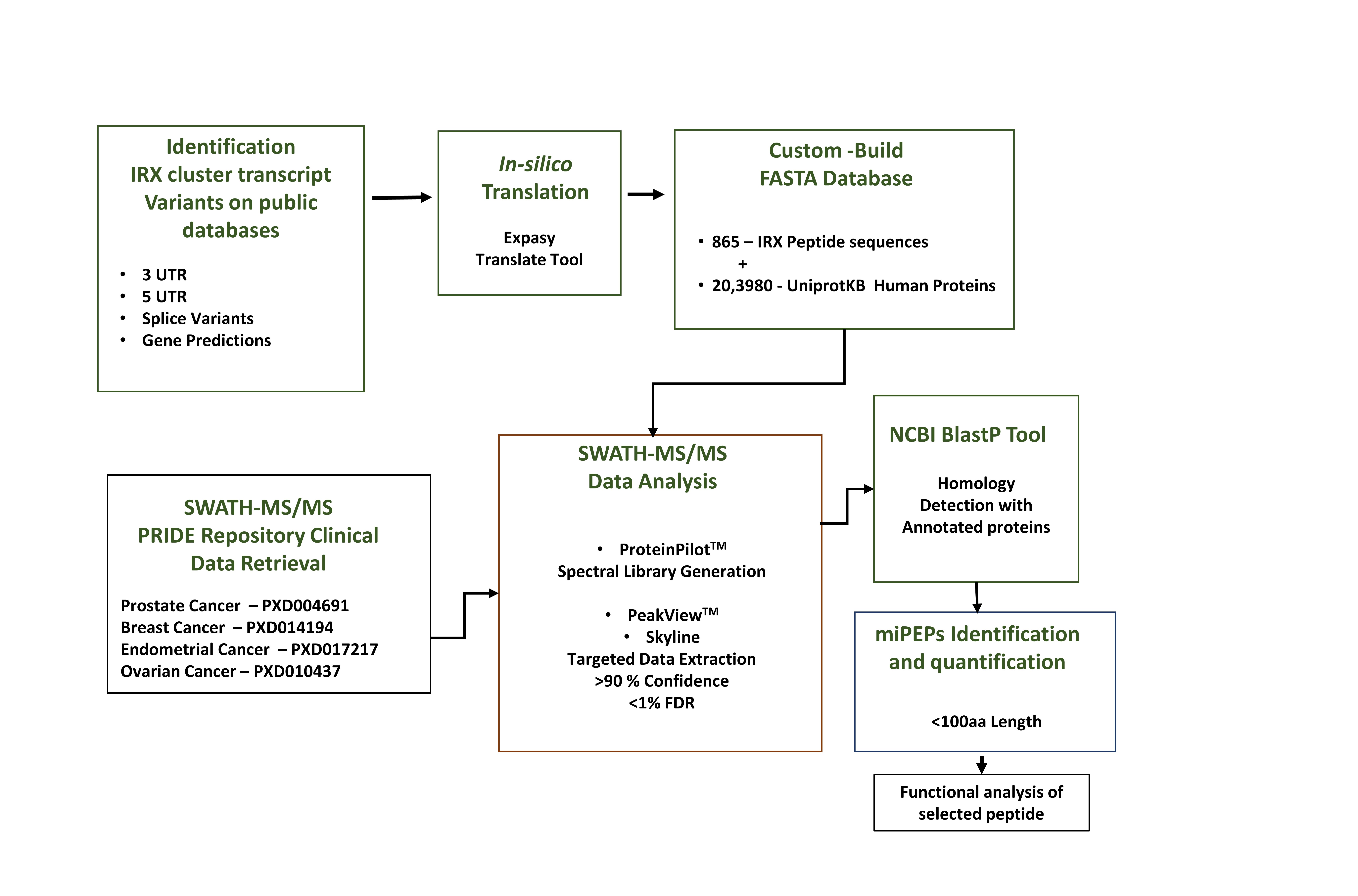

### Supplemental Figure 2

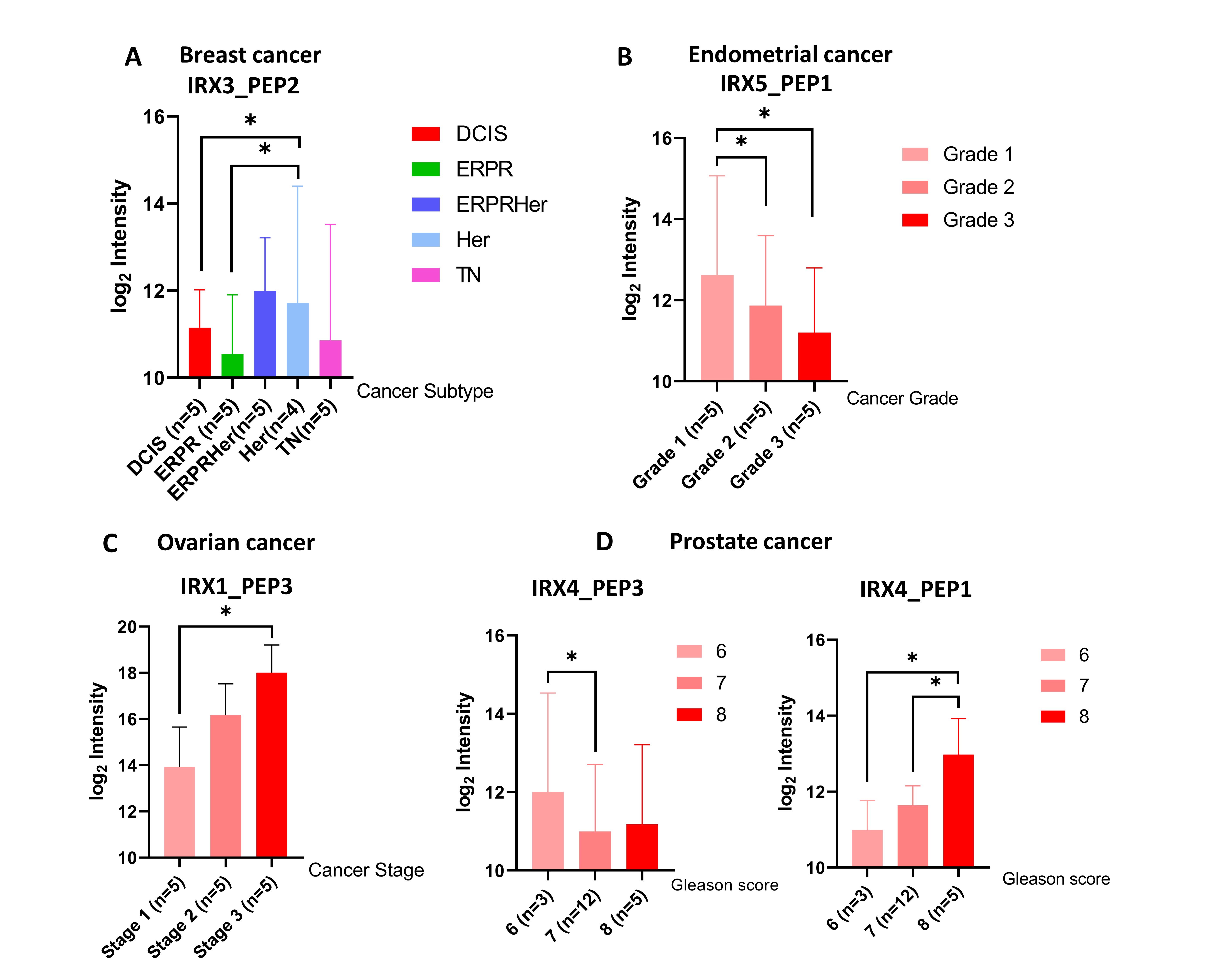

### Supplemental Figure 3

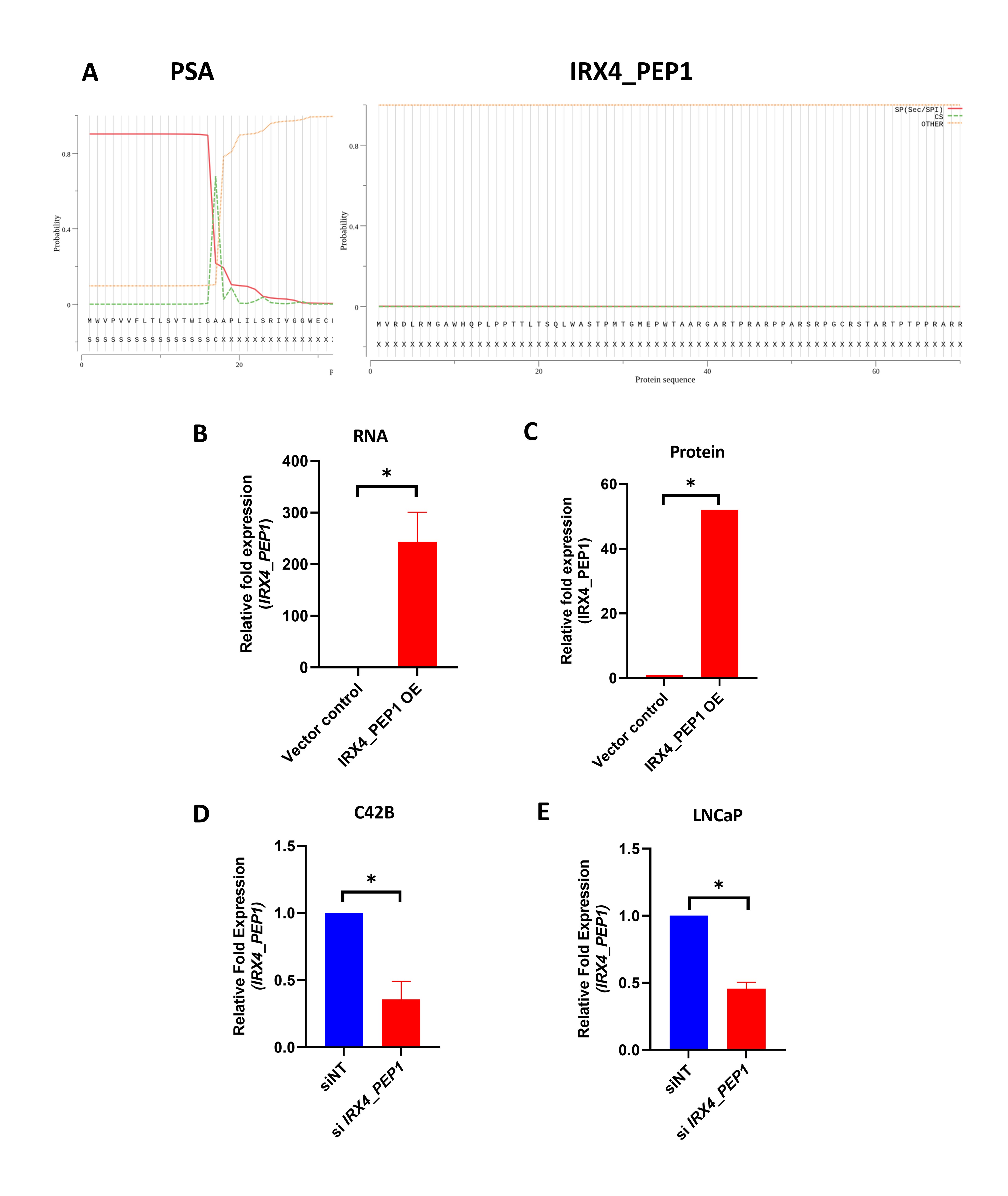

### Supplemental Figure 4

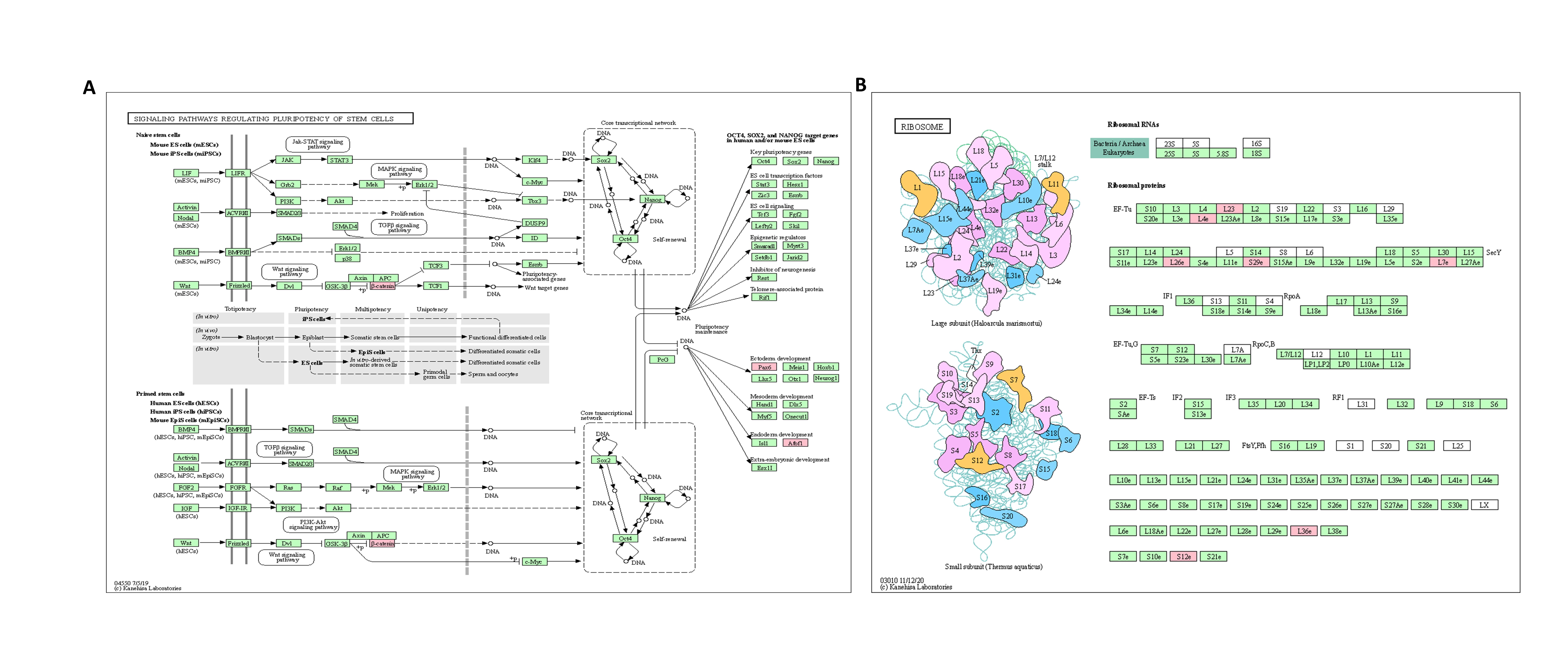
